# Fine-scale animal proximity detection and localization via multi-sensor biologgers

**DOI:** 10.1101/2025.09.09.674951

**Authors:** Will Rogers, Sierra J Mattingly, Jenna E Kohles, Nils Linek, Hannah J Williams, Ivan Lenzi, Georg Wilbs, Marcel Escher, Nina Richter, Louis van Schalkwyk, Vanessa O Ezenwa, Martin Wikelski, Dina KN Dechmann, Timm A Wild

**Affiliations:** Department of Ecology and Evolutionary Biology, Yale University, New Haven, CT, USA; Department of Migration, Max Planck Institute of Animal Behavior, Radolfzell, Germany; Department of Biology, University of Konstanz, Konstanz, Germany; Centre for the Advanced Study of Collective Behaviour, University of Konstanz, Konstanz, Germany; Smithsonian Tropical Research Institute, Panama, Republic of Panama; Office of the State Veterinarian, Department of Agriculture, Land Reform and Rural Development, Kruger National Park, Skukuza, South Africa; Department of Veterinary Tropical Diseases, Faculty of Veterinary Science, University of Pretoria, Pretoria, South Africa

**Keywords:** Proximity logging, biologging, received signal strength indicator (RSSI), trilateration, sensor validation, animal interactions, social behavior

## Abstract

Accurately quantifying spatial interactions is central to understanding social behavior, information flow, predator-prey dynamics, and disease transmission. Proximity loggers that record received signal strength indicator (RSSI) offer a promising approach for estimating pairwise distances, particularly in environments where GPS is unavailable or imprecise. However, RSSI is often dismissed as too noisy for fine-scale inference, with performance that depends on environmental conditions, tag orientation, and between-device variability. Incorporating additional tag-measured data may improve RSSI performance and enable its use as a continuous measure of distance in variable environments.
Here, we assess the utility of continuous RSSI as a fine-scale distance estimator and localization tool using a novel multi-sensor WiFi biologger (WildFi). We conducted four experiments: (1) testing how tag orientation affects RSSI-distance relationships; (2) evaluating whether environmental covariates measured by onboard sensors improve proximity estimates; (3) assessing the accuracy of trilateration-based tag localization using fixed gateway arrays; and (4) comparing RSSI- and GPS-inferred proximity in free-ranging Egyptian fruit bats (*Rousettus aegyptiacus*).
While RSSI alone could predict distance with reasonable accuracy, incorporating additional tag-sensed information (*e.g.*, temperature, humidity, barometric pressure) and accounting for tag-level heterogeneity significantly improved predictive accuracy. Based on RSSI predictions, we could estimate tag location with a median error of 2.6 meters, accurate enough to indirectly estimate proximity networks without tag-to-tag communication. In deployments on free-flying bats, we found that RSSI and GPS were only weakly concordant, with GPS unreliable for detecting fine-scale interactions (<50 m). In contrast, RSSI could capture both fine-scale and some long-range interactions up to ∼250m.
These findings highlight RSSI’s potential as a robust metric for proximity logging, particularly when combined with multi-sensor data and pre-deployment validations. Integrating multi-sensor data streams further enhances RSSI interpretability. Future biologger designs should prioritize synergy among data streams for integrated insights into proximity and animal behavior.

**Data and code for peer review statement:** Data and code to reproduce the results of the paper are provided in a zip folder for peer review. We also provided our compiled code.

## Introduction

Measuring fine-scale interactions between animals is essential for understanding social structure, information exchange, predator-prey dynamics, and disease transmission (Krause et al. 2013, Smith and Pinter-Wollman 2021, Suraci et al. 2022), processes with implications for conservation and management (Snijders et al. 2017). These processes depend not only on whether animals encounter each other, but also how close they are, how often they interact, and for how long—requiring tools that capture proximity at relevant spatial and temporal scales.

Wireless technologies like Bluetooth Low Energy (BLE) and WiFi offer one such solution by recording received signal strength indicator (RSSI), which declines with distance and can serve as a continuous proxy for proximity between animal-borne sensors. While many RSSI applications discretize interactions into binary “contact” or “non-contact” events (Cross et al. 2012), these thresholds can be misaligned with underlying biological processes and obscure important variation in interaction frequency, duration, or intensity (Craft 2015). Treating RSSI as a continuous variable rather than a discrete contact metric may provide more nuanced and biologically interpretable insights into proximity-dependent processes without imposing rigid assumptions about contact definitions (St Clair et al. 2015, Ripperger et al. 2016).

However, RSSI-based distance estimates are sensitive to environmental and technical variation (Drewe et al. 2012, Bettaney et al. 2015, Rutz et al. 2015), such as ambient weather conditions (Marfievici et al. 2013), the habitat structure (Böhm et al. 2009), antenna orientation (Prange et al. 2006), and variation between tags (Boyland et al. 2013). These sources of variation may reduce the precision of RSSI-based estimates, particularly in field conditions. This leaves open questions about whether RSSI can serve as an accurate proxy of distance and how to refine RSSI predictions.

One approach to enhance RSSI performance is to combine them with other tag-sensed information related to RSSI signal error. For example, multi-sensor biologgers are devices that integrate multiple sensing modalities on the same tag, supporting inferences beyond single sensors alone (Williams et al. 2020). Because RSSI is influenced by weather, collecting temperature, humidity, or barometric pressure simultaneously with RSSI could offer pathways to account for these sources of variation, estimating distance as a function of multiple data streams beyond RSSI alone. Multi-sensor approaches remain largely untested in proximity logging – especially in free-ranging animals and under real-world environmental variability – leaving open questions about how beneficial multi-sensor inferences are to RSSI predictions.

If RSSI can reliably estimate distance, it also opens the door to spatial localization by inferring an animal’s position in space by comparing RSSI from a mobile tag to three or more fixed receivers, or “gateways.” This technique, referred to as trilateration, has been widely applied in automated VHF and UHF telemetry systems (Kays et al. 2011, Kluge et al. 2020, Ripperger et al. 2020, Paxton et al. 2022). In settings where GPS is unavailable, imprecise, or power-intensive, such as caves, dense forests, or burrows, RSSI-based localization offers a lightweight alternative. Moreover, if multiple tagged individuals are localized with sufficient accuracy, their relative positions can be used to reconstruct pairwise proximity networks without direct tag-to-tag communication. While promising, the accuracy of RSSI-based localization may be limited by signal variability. Whether integrating additional tag-measured context (*e.g.*, weather or line-of-sight conditions) can improve trilateration performance remains an open question.

As animal-borne GPS tracking continues to expand in both taxonomic breadth and global scale (Kays and Wikelski 2023), there is growing interest in using GPS not only to track movement but also to infer interactions between individuals (He et al. 2023, Yang et al. 2023), relevant to information exchange (Merkle et al. 2024) or disease spread (Wilber et al. 2022).

However, GPS is accompanied by positional error, ranging from several to tens of meters depending on environment (Frair et al. 2010), and variable time-to-fix delays, ranging from several seconds to minutes (He et al. 2023). As a result, GPS often overestimates distance between tags (He et al. 2023), possibly missing important fine-scale interactions. In contrast, proximity sensing may offer time-synchronized observations (Wild et al. 2024) and greater accuracy than GPS at fine spatial scales. Though, empirical comparisons are needed to evaluate when and how these methods yield different conclusions about animal interactions.

In this study, we used the multi-sensor WildFi biologger (Wild et al. 2022b) to evaluate RSSI’s performance as a tool for proximity estimation (both continuous and contact-based) and localization under varying conditions. We conducted three experiments and a field trial: (1) testing how tag orientation affects RSSI-distance relationships (**Fig. 1A**); (2) assessing whether tag-measured temperature data improves proximity estimates (**Fig. 1B**); (3) evaluating trilateration using RSSI in a fixed gateway array (**Fig. 1C**); and (4) comparing GPS- and RSSI-inferred proximity in free-ranging Egyptian fruit bats (*Rousettus aegyptiacus*) (**Fig. 1D**). Across these tests, we asked whether RSSI alone is sufficient for fine-scale inference, and whether incorporating sensor-derived covariates improved accuracy. By combining experimental validation with field deployment, we assessed the value of multi-sensor biologging for proximity inference in behavioral ecology.

**Figure 1:**
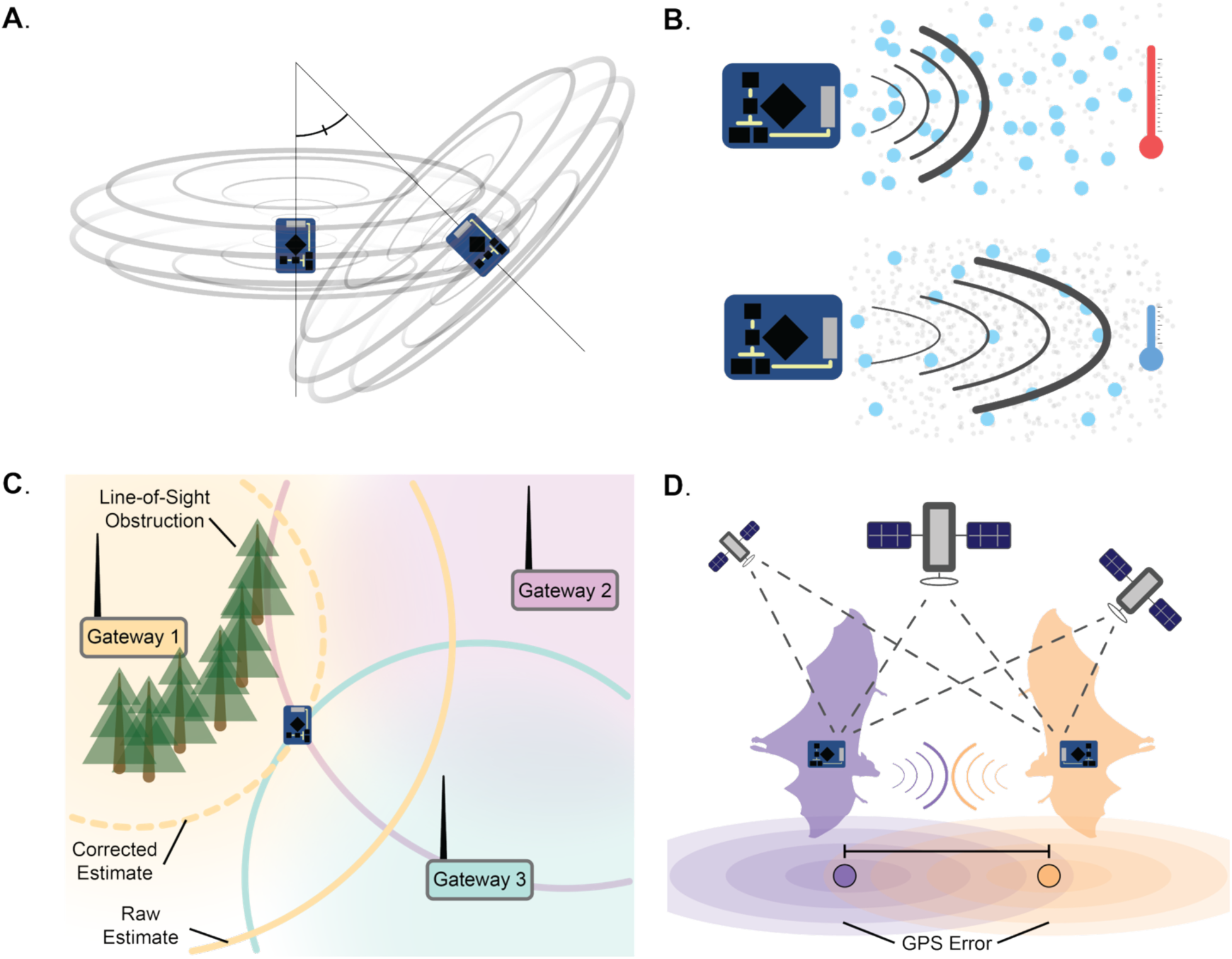
**Challenges and applications in proximity sensing addressed by experiments and field trials**. (A) Differences in tag orientation influence signal propagation, altering the relationship between Received Signal Strength Indicator (RSSI) and true distance. (B) Local weather conditions, such as temperature, humidity, and air pressure, affect air density and refractive properties, causing signal attenuation. (C) Stationary arrays enable signal localization through trilateration, though array design and obstructions may impact accuracy. (D) GPS and RSSI can both be used for proximity sensing, but GPS error may compromise fine-scale proximity estimates.

## Materials and Methods

### Experiment 1: RSSI and tag orientation

RSSI typically decays with distance, but factors such as tag orientation can influence how anisotropic signals propagate through space (Rutz et al. 2015). To test the effect of tag orientation on RSSI-distance relationships, we conducted an experiment with two WildFi tags placed on 1-m-high stakes at distances from 3.5 cm to 10 m, with line-of-sight between tags. We recorded RSSI every minute for 10 minutes at each of five relative orientations: (1) antennae facing together, (2) facing away, (3) perpendicular, (4) collinear in opposite directions, and (5) collinear in the same direction.

### Experiment 2: RSSI and tag-sensed temperature

Signal propagation in WiFi and other UHF systems can vary with ambient environmental conditions, including air temperature (Marfievici et al. 2013). To evaluate how ambient temperature influences RSSI-based distance estimates, we deployed 35 WildFi tags in arrays at known distances (1 cm to 21.4 m) and uniform orientation. Tags were placed 30 cm above the ground on plastic stools and recorded RSSI and onboard temperature every 30 min from 5:00– 20:00 across trials lasting 30 minutes to 2 days. Natural temperature variation (2.1–53.5 °C) enabled assessment of temperature effects on RSSI under field conditions.

### Experiment 3: RSSI and gateway-based trilateration

RSSI can be used to trilaterate mobile tags; however, obstructions can distort RSSI and affect localization accuracy (Rutz et al. 2015). To test whether tag-measured temperature and line-of-sight conditions influence RSSI–distance estimates and localization, we deployed 16 fixed WildFi gateways across six, 10.5 m × 21 m walled enclosures (**Figs S1-2**, approximately 1.3 ha). Tags were placed at known positions (2-15 m apart) on 30 cm-high stools and recorded RSSI and temperature every 30 min from 05:00-20:00. For each tag–gateway pair, we recorded whether a physical wall obstructed the signal to define a binary “line-of-sight” variable.

### Field Trial: RSSI vs. GPS and on-animal inferences

While GPS error is predicted to inflate pairwise distances (He et al. 2023), it is unclear whether RSSI is more accurate in pairwise distance estimation. We compared RSSI- and GPS-based proximity using data on 154 free-ranging Egyptian fruit bats captured in Cyprus, fitted with WildFi biologgers programmed to record RSSI, GPS locations, and environmental data (temperature, relative humidity, and barometric pressure) every 5 min for approximately 5 days. GPS sampling faced variable time-to-fixed intervals, ranging from 1 to 255 seconds (mean = 20.6 s). We minimized the effect of RSSI-GPS asynchrony by only retaining observations with ≤30 s time difference between paired GPS fixes and RSSI measures. Though, thresholds up to 300 s did not alter inferences (**Figs S3–5**).

### Statistical analysis

We modeled tag–tag and tag–gateway distances as a function of RSSI using Gamma regression models with a log-link based on fit comparisons to Gaussian and log-Gaussian alternatives (**Fig. S6**). In addition to RSSI-only models, we fit more complex models incorporating additional variables (tag orientation in **Experiment 1**, temperature in **Experiment 2**, line-of-sight and temperature in **Experiment 3**, and temperature, relative humidity, and barometric pressure in the **Field Trial**). Random effects were included for the tag recording RSSI, and where applicable, for the alternate tag or receiving gateway. All predictor variables were scaled and centered prior to fitting using *glmmTMB* (Brooks et al. 2017), and we assessed model fit using *DHARMa* (Hartig 2018). We evaluated nonlinear RSSI-distance relationships using generalized additive models in *mgcv* (Wood 2010) (**Figs S7-9**). In the **Field Trial**, we found a nonlinear relationship with GPS-inferred distance plateauing at ∼48 m for RSSI values greater than –75 (**Fig. S10**). To address this issue, we took two modeling approaches: first, we log-transformed RSSI to capture the observed nonlinear structure in GPS-inferred distances (**Fig. S11**); second, we alternatively excluded observations with RSSI values > -75, reducing the influence of potential GPS error. Under this filtering approach, the relationship between RSSI and distance more closely resembled the linear exponential decay pattern observed in **Experiments 1–3 (Fig. S12).**

We assessed model performance based on (1) model fit and (2) predictive accuracy on out-of-sample data. First, we compared RSSI-only and more complex model fit using Akaike’s Information Criterion (AIC), with differences >4 indicating strong support for the more complex model (Akaike 1998). We also calculated marginal and conditional *R^2^* of models estimated using *MuMIn* (Barton 2009). Second, we evaluated predictive performance using cross-validations with 25 random splits into training (75%) and testing (25%) data. We generated predictions both conditioned on tag/gateway identity and using only population-level effects (i.e., as if there were no pre-deployment validation data). We evaluated predictive error based on continuous distance predictions (using median *R^2^* and root mean squared error [RMSE]) and accuracy in classifying contact, based on multiple distance thresholds. We compared accuracy against a random classifier determined by the proportion of true “contacts” at various distance thresholds in testing data (*p*), with accuracy equal to *p*^2^+(1-*p*)^2^. We additionally report sensitivity and specificity in **Figs S13-16**.

### Trilateration positional accuracy inferred proximity networks

We tested whether RSSI-based trilateration could accurately localize tags and reconstruct proximity networks within gateway arrays. Using nonlinear least squares (NLS) regression based on previous trilateration approaches (Paxton et al. 2022), we estimated tag positions based on predicted distances to fixed WiFi gateways with known locations. For each tag and time point, we used all gateways observed by a tag (mean = 13.097, range: 10-15) and generated two types of predictions: (1) conditional on tag-specific random effects, or (2) using only population-level effects. To account for the expected increase in error with weaker RSSI, we assigned exponentially decaying weights to tag–gateway distance predictions, and we initialized NLS models with start locations based on the weighted average of all gateway coordinates. We quantified Euclidean error of location estimates, and we compared correlations between trilateration-inferred and true proximity networks.

### RSSI- vs. GPS-inferred proximity and contact networks

We compared RSSI- and GPS-inferred proximity networks using data from free-flying bats. Using predictions from the top **Field Trial** model (based on a dataset filtered to RSSI ≤ - 75), we constructed (1) continuous networks based on average pairwise distances, and (2) binary contact networks at thresholds from 0.5-500 m. We generated equivalent networks from GPS by computing pairwise distances for GPS fixes. We compared the correlation between respective distance and contact networks using Mantel tests in *vegan* (Oksanen et al. 2013).

We assessed whether discrepancies between RSSI and GPS were driven by GPS error by simulating positional error from gamma distributions fit to empirical GPS data (Wild et al. 2022a) (**Figs S13-19)**. We generated 1,000 replicate GPS position pairs for true distances ranging from 0.01-1,000 m, incorporating stochastic error into each fix, and calculated apparent inter-tag distances. We compared simulated GPS distances to empirical RSSI estimates and calculated percent error across distances. This allowed us to assess whether observed divergence between GPS and RSSI proximity was consistent with expected GPS error and how it compared to known RSSI error from **Experiments 1-3**. We performed all analyses using Program R version 4.3.2 (Team 2021).

### Animal Handling

Data from Egyptian fruit bats were collected with ethical permission from the Ministry of Agriculture, Agricultural Development and Environment, Cyprus, under permit numbers 02.15.001.003/04.05.002.005.006/02.15.007.003.002 (10/10/2022).

## Results

### Experiment 1: RSSI and tag orientation

We first tested whether tag orientation affected the relationship between RSSI and distance. Using a controlled setup with two tags spaced from 3.5 cm to 10 m, we found that tag orientation significantly influenced RSSI across all distances (likelihood ratio test [LRT] *χ^2^* = 1859, df = 4, *p*-value < 0.001; **Fig. 2A**, **Table 1**). Full models including orientation were favored over RSSI-only models based on AIC comparisons, and full models explained more variance in observed data (conditional *R^2^* = 0.963, marginal *R^2^* = 0.963) than RSSI-only models (conditional *R^2^* = 0.800, marginal *R^2^* = 0.800, **Table 1**). Even without information about orientation, RSSI alone was a strong predictor of tag-tag proximity (*β* = -1.746, standard error [SE] = 0.031; LRT *χ^2^* = 3227, df = 1, *p*-value < 0.001, **Table 1**).

**Figure 2:**
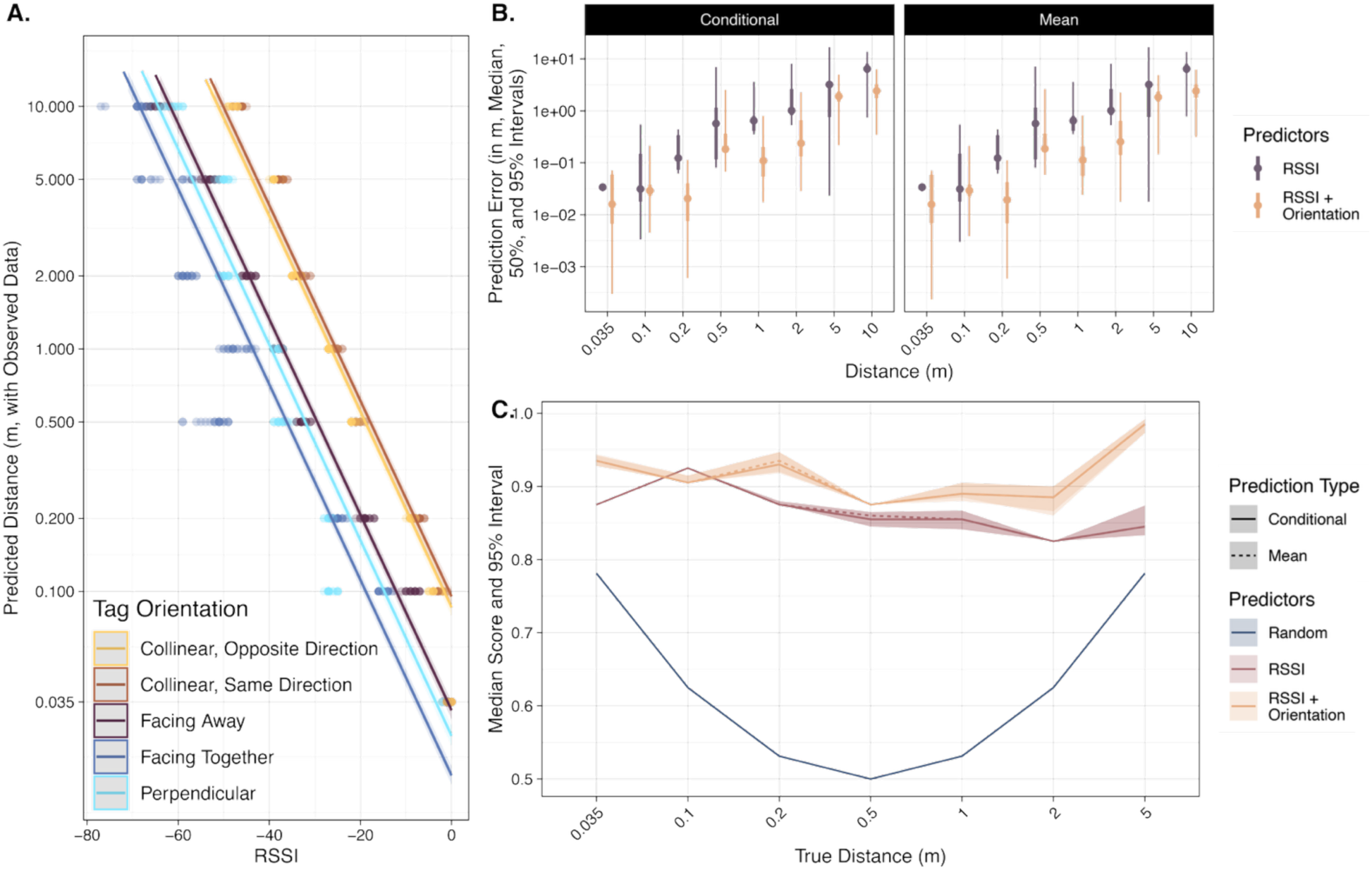
Effect of orientation on RSSI performance in Experiment 1. (A) RSSI-distance relationships differ based on tag orientation (color). Shown are observed data, model estimates, 95% confidence intervals. (B) RSSI predictive error is generally lower for predictions including orientation, and error increases with pairwise distance. Shown are median, 50%, and 95% intervals of out-of-sample prediction error from cross-validations. Facets compare predictions conditioned on tag-level random effects and versus population-level average predictions. (C) Median and 95% intervals for model accuracy in classifying tag-tag communications at or below various true distance thresholds. Solid lines represent metrics based on predictions incorporating tag-level random effects, while dashed lines reflect population-only main effects. Metrics for a random classifier (blue) are also included for comparison, derived from the proportion of true contacts at each threshold.

**Table 1:** Summaries and comparison of generalized linear mixed-effect models of known distance based on RSSI and orientation from **Experiment 1** with Gamma log-link function (*n_tags_* = 2, *n_observations_* = 800). Estimates are on link scale with standard error in parentheses. Stars denote p-values with: * p-value ≤ 0.05, ** p-value ≤ 0.01, *** p-value ≤ 0.001.

|  | Null | Orientation |
| --- | --- | --- |
| Intercept | -0.107<br>(0.027)*** | 0.418<br>(0.038)*** |
| RSSI | -1.746<br>(0.031)*** | -1.891<br>(0.016)*** |
| Collinear, Same Direction |  | 0.103<br>(0.049)* |
| Facing Away |  | -0.978<br>(0.049)*** |
| Facing Together |  | -1.597<br>(0.049)*** |
| Perpendicular |  | -1.221<br>(0.05)*** |
| Focal ID Random Effect SD | 1.29e-05 | 2.07e-02 |
| $\Delta AIC$ | 968.1028 | 0 |
| Conditional $R^2$ | 0.800 | 0.936 |
| Marginal $R^2$ | 0.800 | 0.936 |

In cross-validations, full models including orientation (median *R^2^* = 0.833, median RMSE = 1.36 m) outperformed RSSI-only models (median *R^2^* = 0.503, median RMSE = 3.87 m, **Table S1**). Continuous prediction error tended to increase proportionately with true distance, but orientation-informed models maintained lower error throughout yielding a median error of 11 cm at 1 m, 1.90 m at 5 m, and 2.43 m at 10 m (**Fig. 2B)**. Classifications of contact based on different distance thresholds also favored full models including orientation over RSSI only models (**Fig. 2C, Fig. S13**). Yet, both RSSI-only and full models performed better than a random classifier with median accuracy greater than 82.5% and 87.5% across all distance thresholds, respectively (**Fig. 2C**). These results confirm that tag orientation introduces meaningful signal variation, but that RSSI alone can still yield reasonably accurate proximity estimates in the face of variation in tag orientations.

### Experiment 2: RSSI and tag-sensed temperature

We next tested whether tag-sensed ambient temperature influenced the relationship between RSSI and distance. Using 25 tags spaced from 1 cm to 21.5 m, we found that temperature had a significant effect on predicted distance, accounting for RSSI (*β* = -0.178, SE = 0.001, LRT *χ^2^* = 32801, df = 1, *p*-value < 0.001, **Fig. 3A**, **Table 2**). Models including temperature were also favored over RSSI-only models based on AIC comparisons and explained more variation in the observed data (conditional *R^2^* = 0.836, marginal *R^2^* = 0.728) than RSSI-only models (conditional *R^2^*= 0.814, marginal *R^2^* = 0.704, **Table 2**). Tag-level random effects were important in RSSI variation, explaining approximately 10–11% of variation. Notably, RSSI alone was a strong predictor of pairwise distances, even when ambient weather conditions were unaccounted for (*β* = -1.093, SE = 0.001, LRT *χ^2^* = 1035446, df = 1, *p*-value < 0.001).

**Figure 3:**
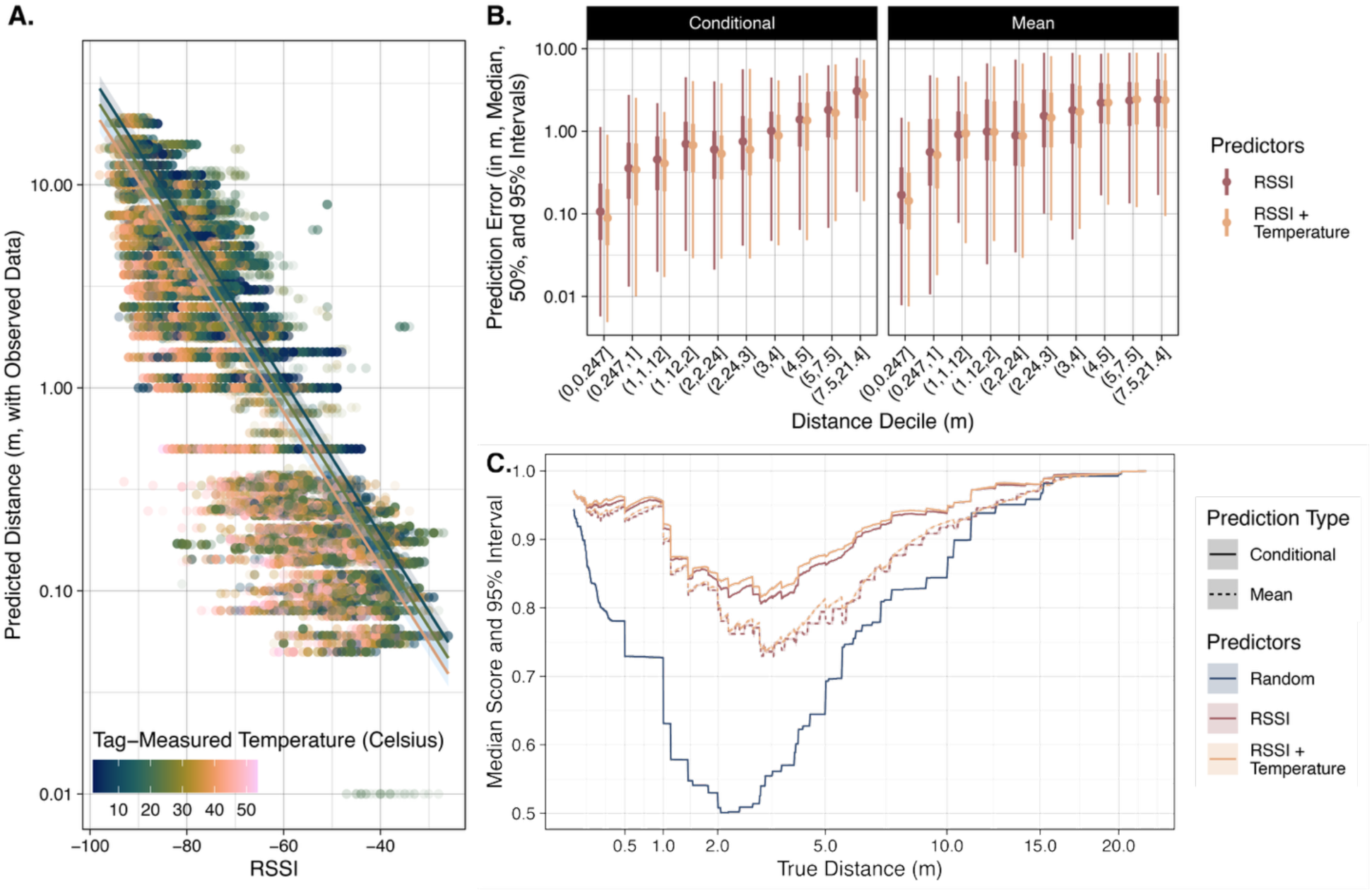
Effect of weather conditions on RSSI performance in Experiment 2. (A) RSSI-distance relationships differ based on tag-measured temperature (color). Shown are observed data, model estimates, 95% confidence intervals. (B) RSSI predictive error is slightly lower for predictions including tag-measured temperature, and error increases with pairwise distance. Shown are median, 50%, and 95% intervals of out-of-sample prediction error from cross-validations. Facets compare predictions conditioned on tag-level random effects and versus population-level average predictions. (C) Median and 95% intervals for model accuracy in classifying tag-tag communications at or below various true distance thresholds. Solid lines represent metrics based on predictions incorporating tag-level random effects, while dashed lines reflect population-only main effects. Metrics for a random classifier (blue) are also included for comparison, derived from the proportion of true contacts at each threshold.

**Table 2:**
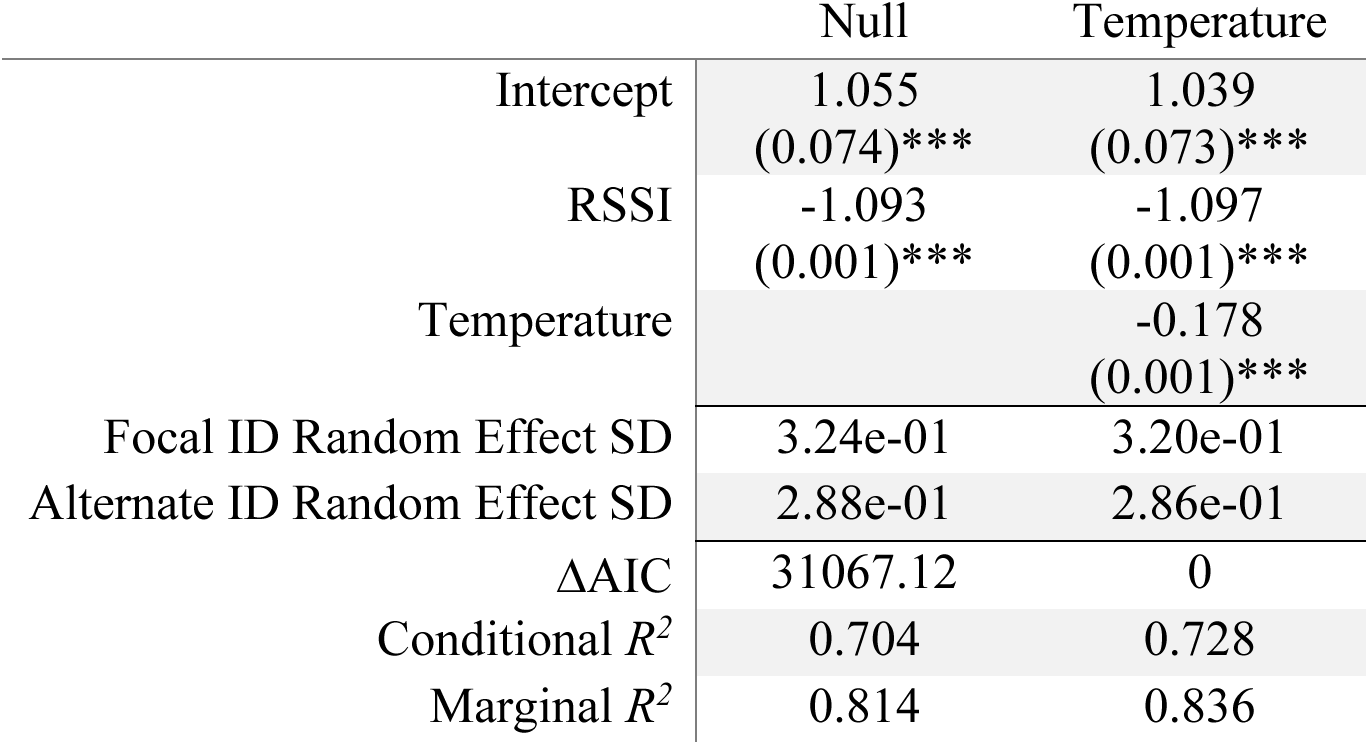
Summaries and comparison of generalized linear mixed-effect models of known distance based on RSSI and tag-measured temperature from **Experiment 2** with Gamma log-link function (*n_tags_* = 35, *n_observations_* = 269,443). Estimates are on link scale with standard error in parentheses. Stars denote p-values with: * p-value ≤ 0.05, ** p-value ≤ 0.01, *** p-value ≤ 0.001.

Cross-validations confirmed that models including temperature produced more accurate predictions (median *R^2^* = 0.697, median RMSE = 1.894 m) than RSSI-only models (median *R^2^* = 0.585, median RMSE = 2.721 m, **Table S2**). Prediction errors increased with true distance: for example, the top model produced median error of 34.2 cm at distances <1 m, 59.9 cm at 2.2-3 m, and 1.67 m at 5-7.5 m (**Fig. 3B**). Contact classification also favored models including temperature over RSSI-only models, but all models outperformed a random classifier (**Fig. 3C, Fig. S14**). Conditioned on tag identity, contact classification accuracy was >80.6% across all distance thresholds; in contrast, predictions assuming population-level averages was only >72.9% accuracy. Fine-scale contacts were exceptionally accurate with approximately 95% accuracy in defining contacts <1 m (**Fig. 3C**). These results demonstrate that RSSI can provide effective proximity estimates, and that accounting for tag-sensed temperature or tag-level heterogeneity can further improve performance.

### Experiment 3: RSSI and tag-gateway communications

We then tested whether tag-measured temperature or information about gateway positioning (specifically, line-of-sight conditions) could increase the predictive performance of RSSI in tag-gateway distance estimates. Across 16 fixed gateways and a range of tag positions within a 64 m × 21 m enclosure, we found that tag-measured temperature affected distance estimates (*β* = -0.022, SE = 0.001, LRT *χ^2^* = 58.3, df = 1, *p*-value < 0.001), and that line-of-sight obstructions affected the relationship between RSSI and true distance (LRT *χ^2^* = 1187.8, df = 1, *p*-value < 0.001, **Fig. 4A**, **Table 3**). Models included both tag-measured temperature and an interaction between RSSI and line-of-sight were favored over RSSI-only models in AIC comparisons and explained substantially more variation in distance estimates (conditional *R^2^*= 0.850, marginal *R^2^* = 0.692) than RSSI-only models (conditional *R^2^* = 0.771, marginal *R^2^* = 0.567, **Table 3**). Tag and gateway random effects explaining approximately 15-20% of variation in distance predictions, indicating a significant role of device variation. When line-of-sight conditions or tag-measured temperature were unaccounted for, RSSI alone remained a strong predictor of distance (*β* = -0.490, SE = 0.006, 95% CI; LRT *χ^2^* = 5955, df = 1, *p*-value < 0.001).

**Figure 4:**
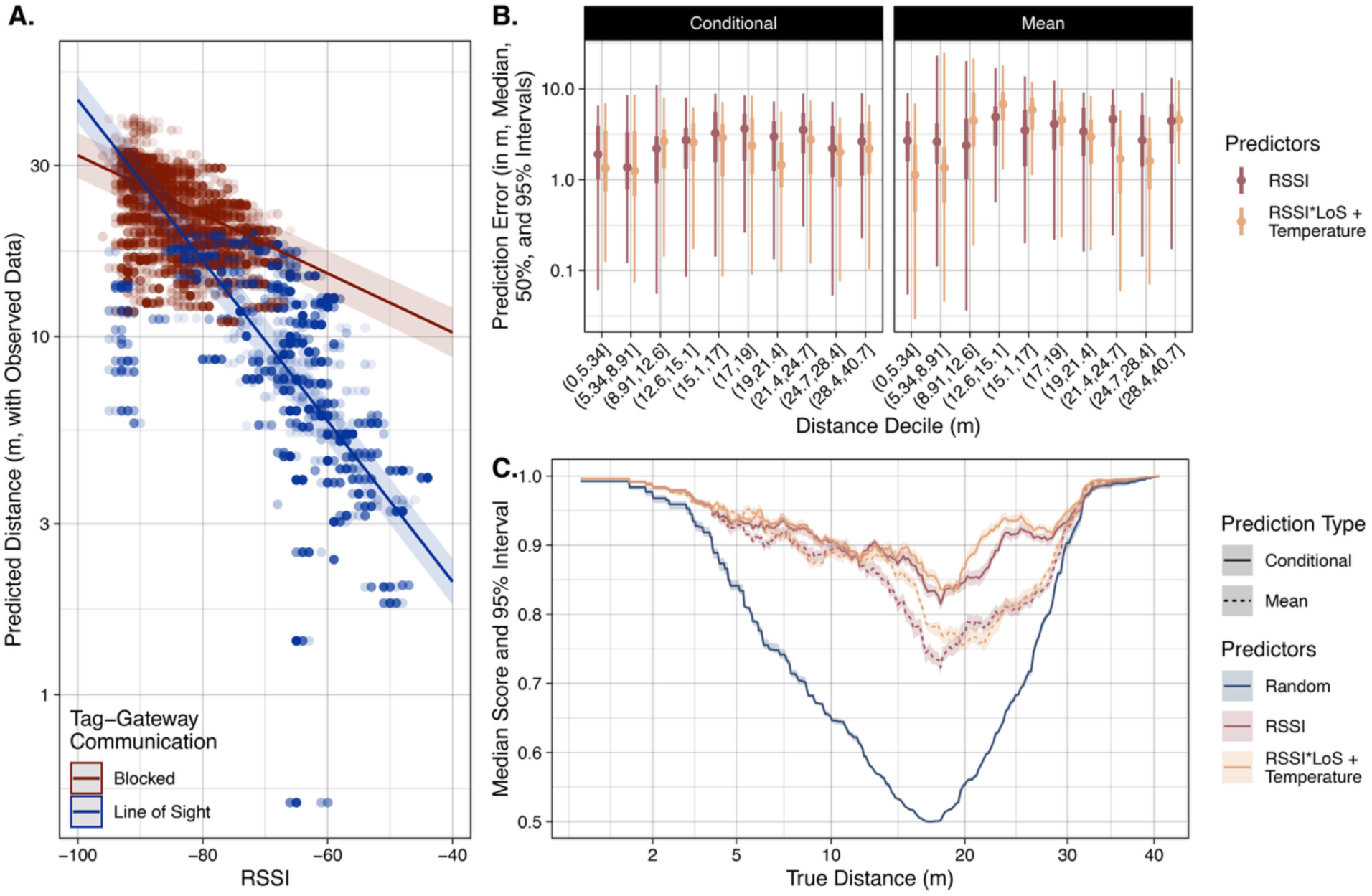
Effect of gateway array design on RSSI performance in Experiment 3. (A) RSSI-distance relationships differ based on line-of-sight conditions (color). Shown are observed data, model estimates, 95% confidence intervals. (B) RSSI predictive error is generally lower for predictions including RSSI-by-line-of-sight (LoS) interactions and tag-measured temperature. Shown are median, 50%, and 95% intervals of out-of-sample prediction error from cross-validations. Facets compare predictions conditioned on tag-level random effects and versus population-level average predictions. (C) Median and 95% intervals for model accuracy in classifying tag-tag communications at or below various true distance thresholds. Solid lines represent metrics based on predictions incorporating tag-level random effects, while dashed lines reflect population-only main effects. Metrics for a random classifier (blue) are also included for comparison, derived from the proportion of true contacts at each threshold.

**Table 3:**
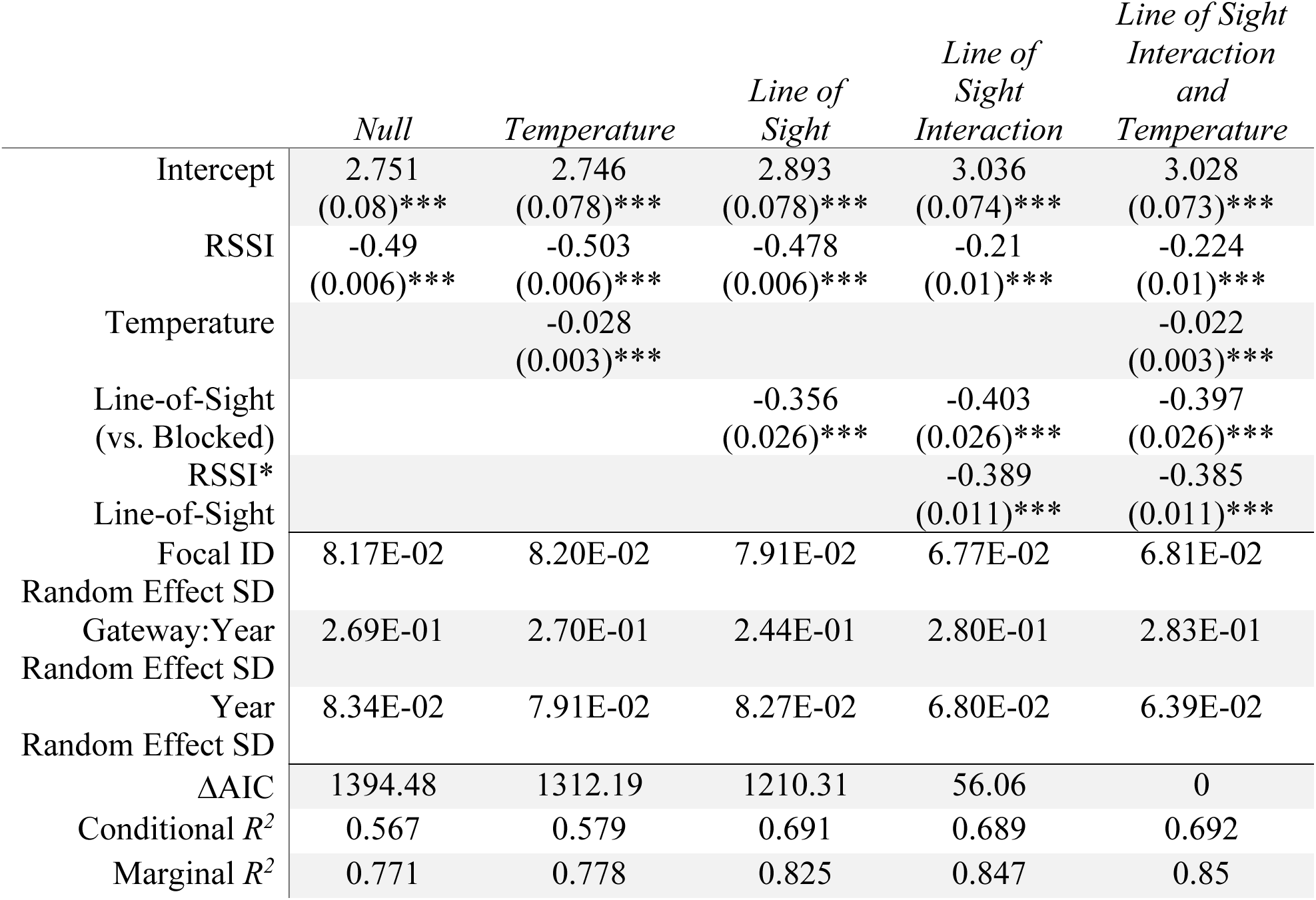
Summaries and comparison of generalized linear mixed-effect models of known distance based on RSSI and tag-measured temperature and line-of-sight conditions from **Experiment 3** with Gamma log-link function (*n_tags_* = 17, *n_gateways_* = 16, *n_observations_* = 10,732). Estimates are on link scale with standard error in parentheses. Stars denote p-values with: * p-value ≤ 0.05, ** p-value ≤ 0.01, *** p-value ≤ 0.001.

In out-of-sample predictions, the full model yielded more accurate distance estimates (median *R^2^* = 0.838, median RMSE = 3.28 m) than RSSI-only models (median *R^2^* = 0.794, median RMSE = 3.78 m, **Table S3**). Prediction error was relatively small compared to true distance, with the top model producing median error of 1.25 m between 5.34 - 8.91 m, 2.36 m between 17 - 19 m, and 2.00 m between 24.7 - 28.4 m. Full models also classified contacts more accurately than RSSI-only models, but all models outperformed a random classifier with greater than 73.3-83% accuracy across all distances (**Fig. 4C**, **Fig. S15**).

### RSSI-Based trilateration using gateway arrays

We next evaluated whether RSSI-based distance estimates could be used to localize tags via trilateration. Using nonlinear least squares trilateration, we estimated a median positional error of 2.59 m (95% interval: 0.560 – 6.04 m) for the full model (including tag-measured temperature and an interaction between RSSI and line-of-sight), and 3.02 m (95% interval: 0.588 – 7.08 m) for the RSSI-only model, conditional on tag-level random effects (**Fig. 5A**, **Fig. S17**). When using population-level predictions, we found roughly similar error for the full model (median = 2.56 m; 95% interval: 0.558 – 6.56 m) but slightly higher error for the RSSI-only model (median = 3.29 m; 95% interval: 0.836 – 7.03 m, **Fig. 5A**).

**Figure 5:**
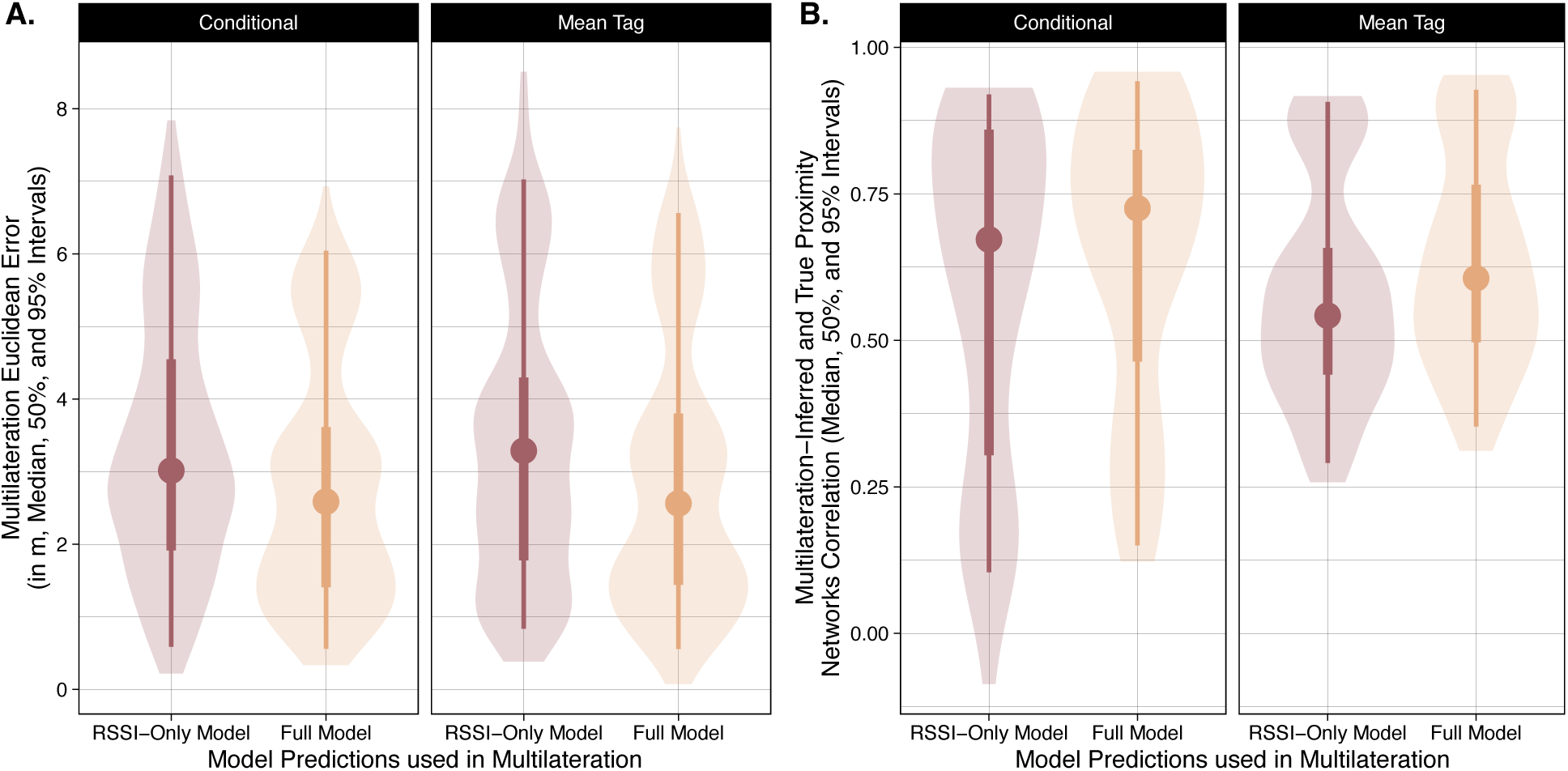
RSSI-based trilateration error in Experiment 3. (A) Euclidean error in trilateration-inferred tag locations within gateway arrays compared to true locations, using an RSSI-only model and a full model incorporating RSSI, line-of-sight conditions, gateway positioning, and tag-measured temperature. (B) Correlation between trilateration-inferred and true pairwise distance networks, comparing trilateration based on RSSI-only versus full model predictions. Facets compare predictions conditioned on tag-level random effects and versus population-level average predictions.

All trilateration-based networks were strongly correlated with the ground-truth proximity networks, though the strength of this correlation varied by model structure. Networks derived from the full model had slightly higher correlations (median = 0.725, 95% interval = 0.150 – 0.942) than those based on RSSI alone (median = 0.672, 95% interval = 0.104 – 0.920), when conditioned on tag identity (**Fig. 5B**, **Fig. S18**). Removing tag-level random effects slightly reduced correlations for both the more complex model (median = 0.606, 95% interval = 0.353 – 0.928) and the RSSI-only model (median = 0.542, 95% interval = 0.291 – 0.907, **Fig. 5B**). These results suggest that RSSI-based inferences in gateway arrays can provide reasonably accurate tag locations useful in generating indirect pairwise proximity estimates.

### Field Trial: RSSI vs. GPS and on-animal inferences

Finally, we sought to understand how RSSI-based distance predictions compared to GPS-based proximity. Using paired RSSI and GPS data from 154 free-flying Egyptian fruit bats, we found that RSSI was associated with GPS-inferred pairwise distances both when RSSI was log-transformed to address non-linearity in the relationship between RSSI-GPS distance (*β* = -0.549, SE = 0.016, LRT *χ^2^* = 1113.8, df = 1, *p*-value < 0.001, **Table 4**), as well as when the dataset was filtered to RSSI-GPS observations less likely to be affected by GPS positional error (*β* = -0.375, SE = 0.019, LRT *χ^2^* = 399.71, df = 1, *p*-value < 0.001, **Table 5**). Under both modeling approaches, we found that models including RSSI and tag-measured temperature, relative humidity, and atmospheric pressure were favored over RSSI-only models (**Fig. 6A**, **Tables 4-5**).

**Figure 6:**
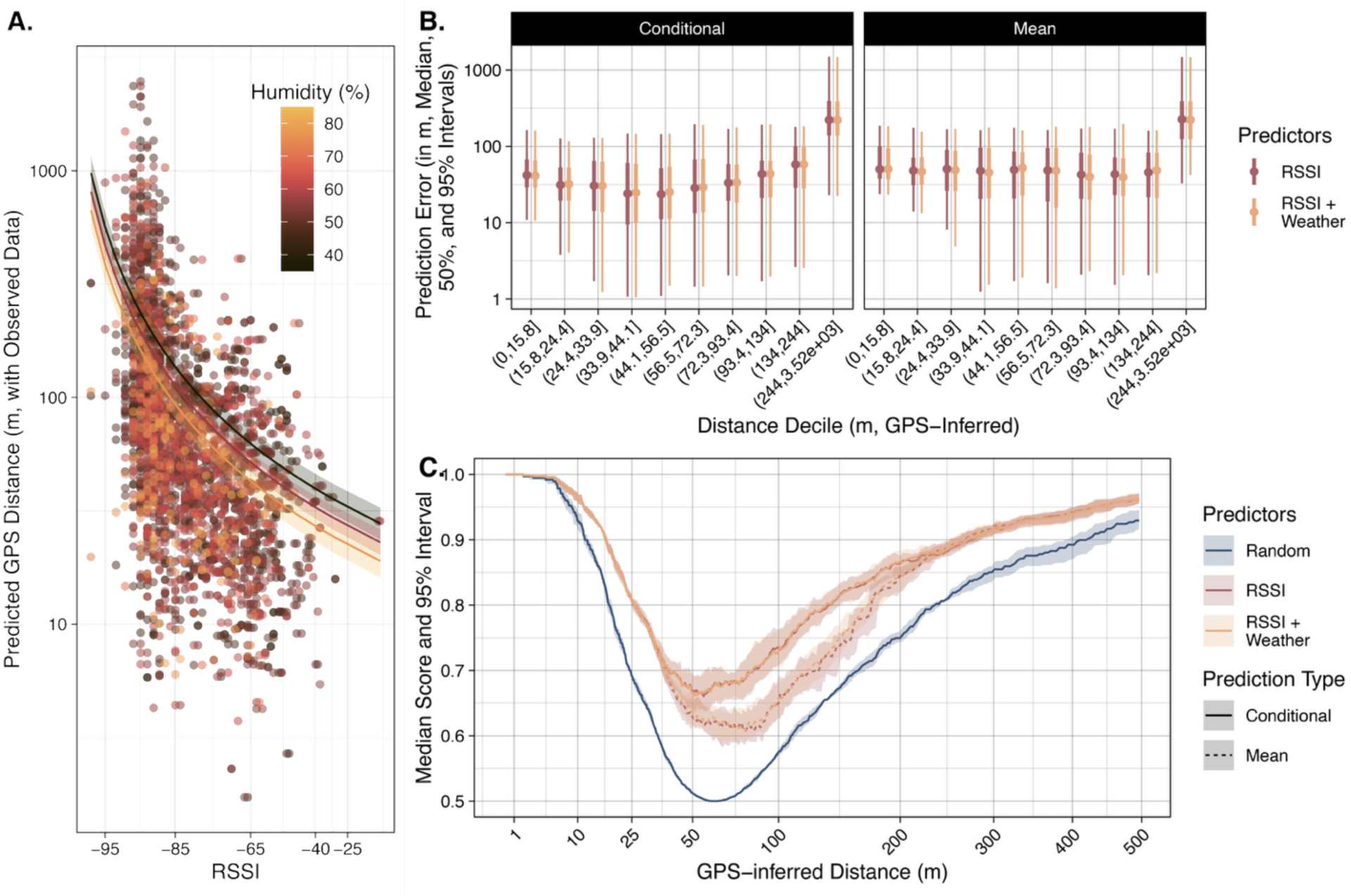
Effect of weather conditions on RSSI performance in the Field Trial. (A) RSSI-distance relationships differ based on relative humidity (color). Shown are observed data, model estimates, 95% confidence intervals. (B) RSSI predictive error is generally lower for predictions including tag-measured weather data (temperature, humidity, and pressure). Shown are median, 50%, and 95% intervals of out-of-sample prediction error from cross-validations. Facets compare predictions conditioned on tag-level random effects and versus population-level average predictions. (C) Median and 95% intervals for model accuracy in classifying tag-tag communications at or below various true distance thresholds. Solid lines represent metrics based on predictions incorporating tag-level random effects, while dashed lines reflect population-only main effects. Metrics for a random classifier (blue) are also included for comparison, derived from the proportion of true contacts at each threshold.

**Table 4:**
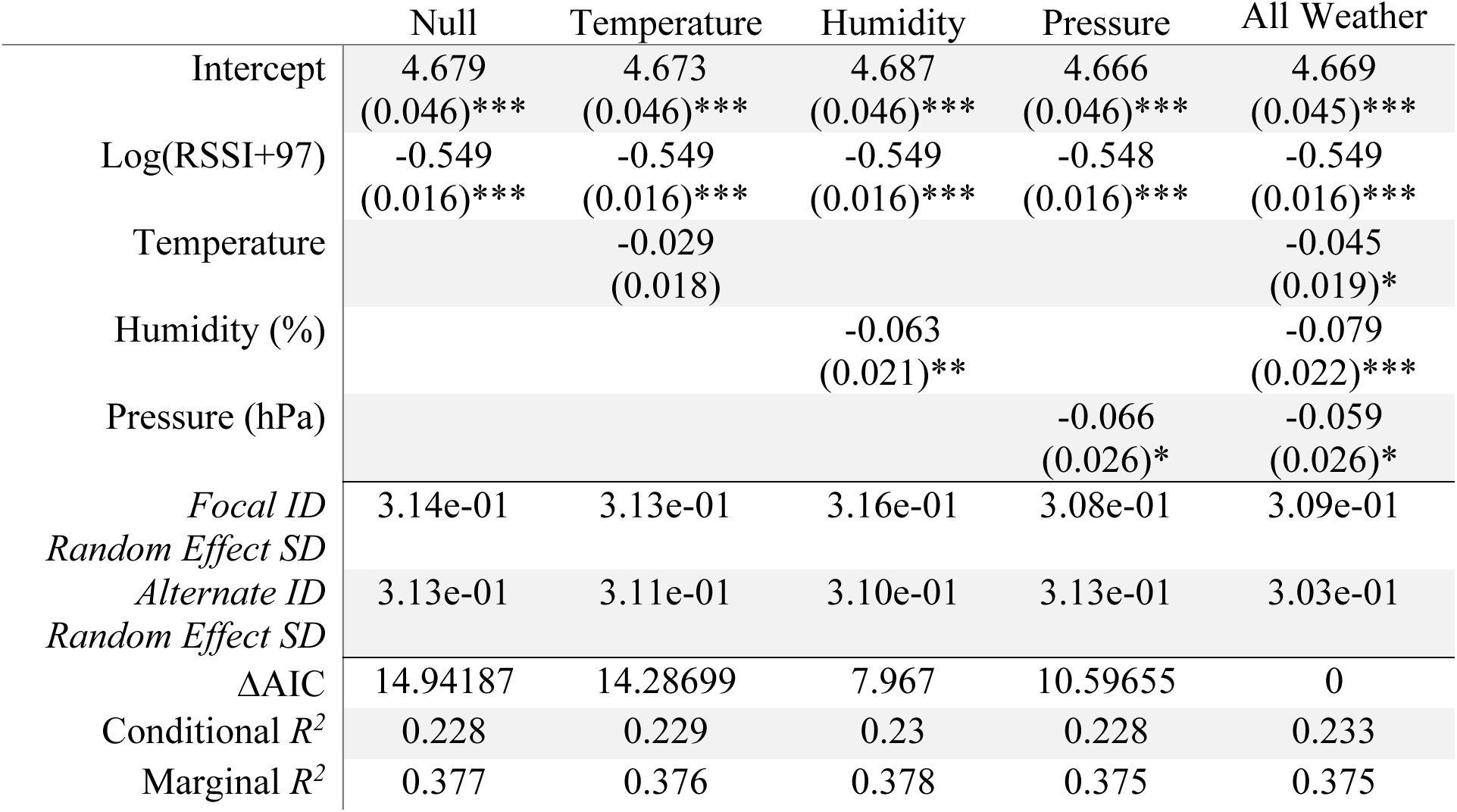
Trial with Gamma log-link function and log-transformed RSSI. Summaries and comparison of generalized linear mixed-effect models of GPS-inferred distance based on RSSI and tag-measured weather data from the **Field Trial** on with Gamma log-link function (*n_tags_* = 136, *n_gateways_* = 136, *n_observations_* = 2,948). Estimates are on link scale with standard error in parentheses. Stars denote p-values with: * p-value ≤ 0.05, ** p-value ≤ 0.01, *** p-value ≤ 0.001.

**Table 5:**
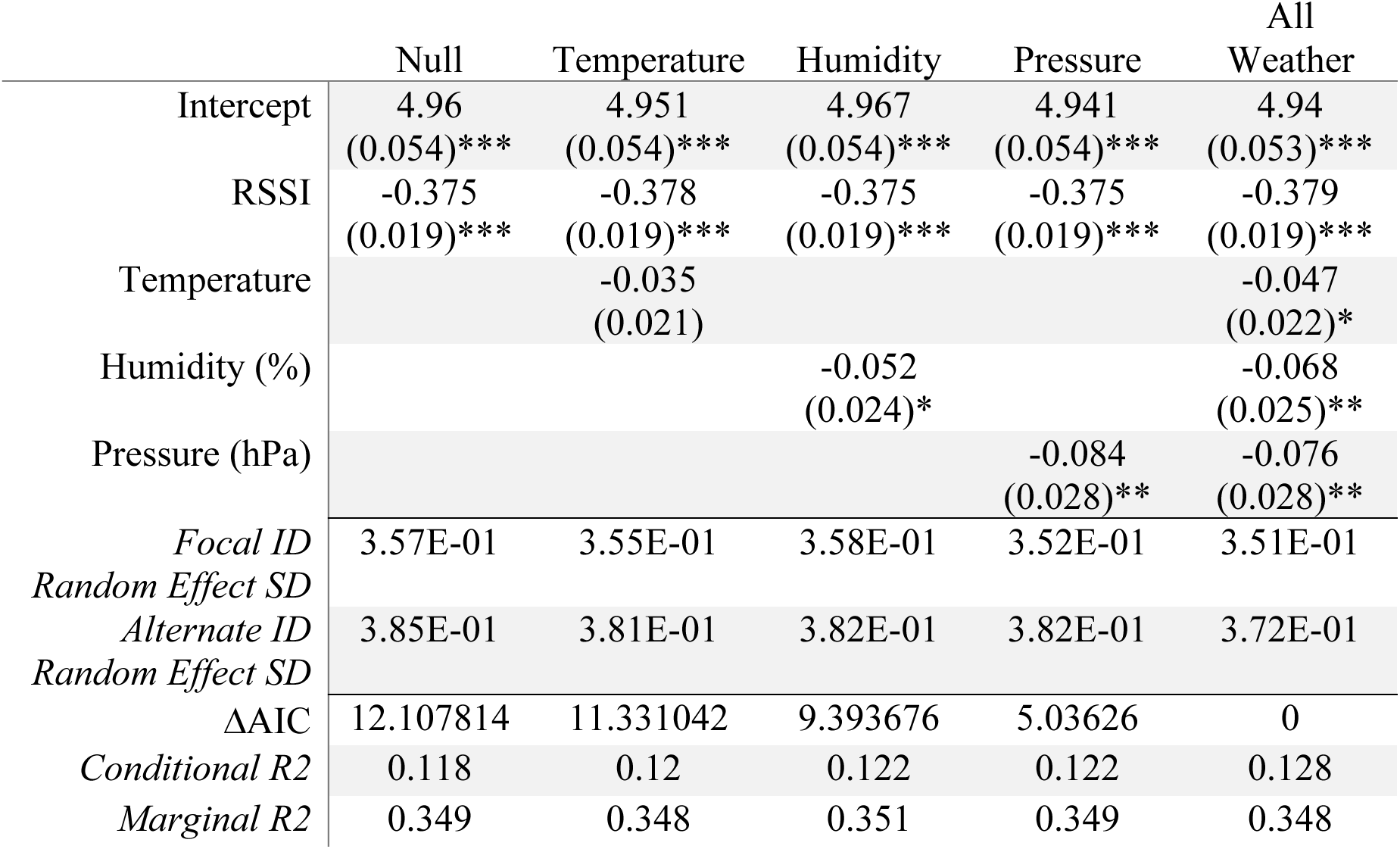
Summaries and comparison of generalized linear mixed-effect models of GPS-inferred distance based on RSSI and tag-measured weather data from the **Field Trial** on with Gamma log-link function, removing observations with RSSI >-75 (*n_tags_* = 134, *n_gateways_* = 134, *n_observations_* = 2,144). Estimates are on link scale with standard error in parentheses. Stars denote p-values with: * p-value ≤ 0.05, ** p-value ≤ 0.01, *** p-value ≤ 0.001.

Cross-validations using all paired RSSI and GPS data showed that including additional tag-measured weather data generated slightly more accurate out-of-sample predictions (median *R^2^* = 0.239, median RMSE = 169.280 m) than RSSI-only models (median *R^2^* = 0.232, median RMSE = 169.415 m, **Table S4**). Distinct from **Experiments 1-3**, prediction error decreased with GPS-inferred distances until approximately 45 m but increased gradually for distances beyond 45 m (**Fig. 6B**). In the top model, for example, median differences between RSSI- and GPS-inferred distances were 41.2 m, 24.7 m, and 58.0 m for GPS-inferred distance bins of 0-15.8 m, 33.9-44.1 m, and 134-244 m, respectively. Full models including tag-measured weather classified GPS-inferred contacts with similar accuracy to RSSI-only models (**Fig. 6B**, **Fig. S16**). While models generally outperformed a random classifier, accuracy was much lower than in **Experiments 1-3** (**Fig. 6C**).

### RSSI- vs. GPS-inferred proximity and contact networks

Using a model excluding observations with RSSI >-75 (where GPS positional error may have been large relative to true pairwise distances), we compared GPS- and RSSI-inferred proximity networks. Across all distances, RSSI- and GPS-inferred average pairwise distance networks were correlated (Mantel statistic *r* = 0.603, *p*-value = 0.001, **Fig. 7A**). However, at close range (<100 m), this relationship disappeared (Mantel statistic *r* = 0.296, *p*-value = 0.135), suggesting divergence in fine-scale proximity estimates (**Fig. 7C**). In total contact networks, we found that RSSI and GPS inferences were unrelated for contacts definitions <5 m, weakly correlated for contacts definitions between 5-25 m, and relatively strongly correlated for contact definitions >50 m (**Fig. 7D**). For example, RSSI- and GPS-inferred 5 m contact networks are largely uncorrelated with each other (**Fig. 7B**).

**Figure 7:**
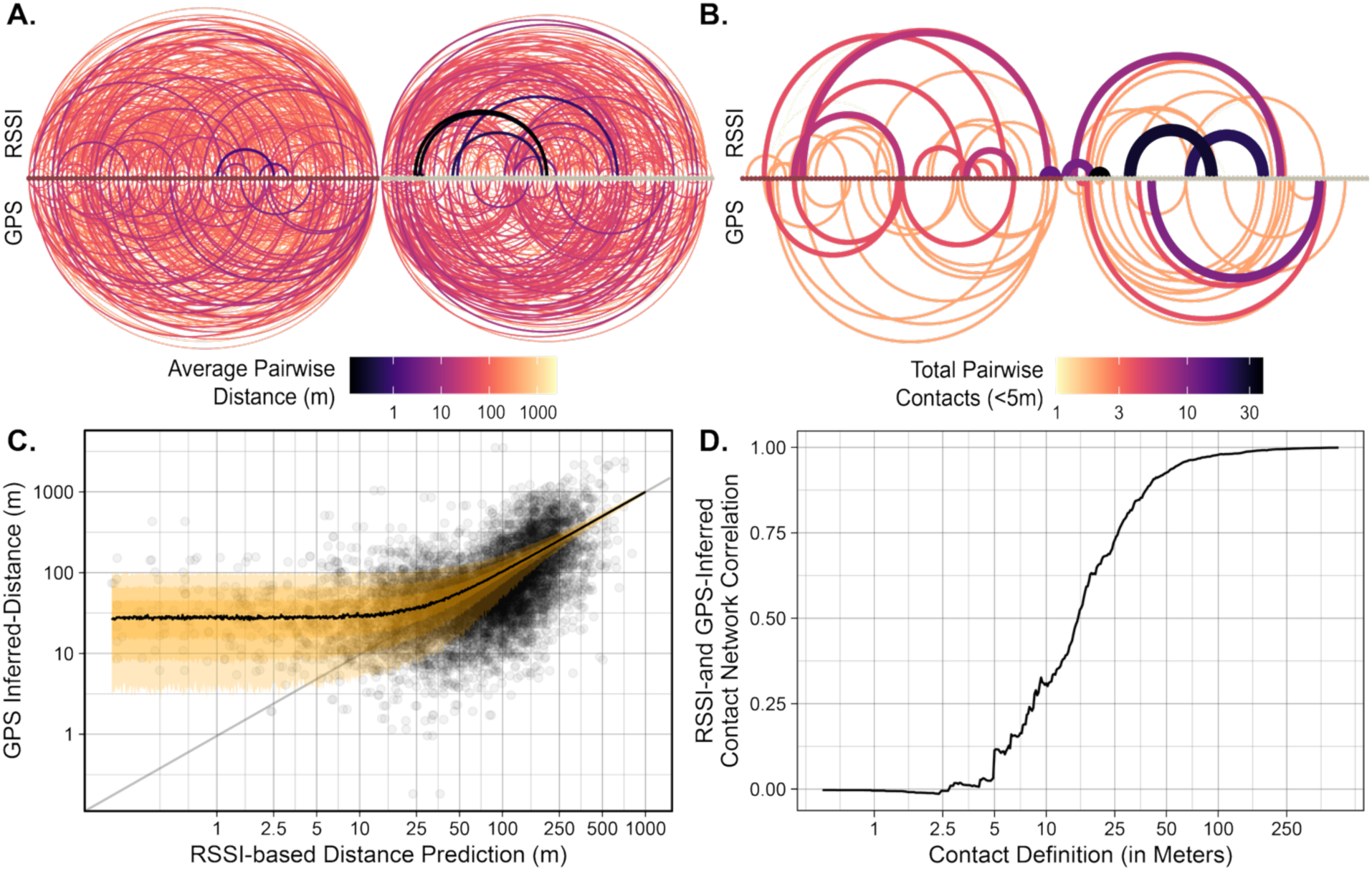
(A) GPS- and RSSI-inferred average pairwise distance (line color and weight) and (B) total contact (<5 m, line color and weight) networks describe different patterns of spatial interactions, particularly at closer pairwise distances. RSSI predictions are conditioned on tag random effects, tag-measured weather data. Nodes within networks are individual bats tagged in 2023 (left, red) or 2024 (right, grey). (C) The disparity in pairwise distance from RSSI predictions and GPS observations is most apparent at distances <50 m (points). Simulations using empirical GPS error recover the observed pattern of inflated pairwise distance estimates (median, 50%, 85%, and 95% intervals shown by black line and shaded orange regions). (D) Similarly, the correlation between RSSI and GPS total contact network declines for contact definitions <50 m, and contact networks are largely uncorrelated for contacts <5 m.

Simulated GPS data, generated using a gamma distribution fit to empirical chip error (**Figs S19-21**), showed consistent overestimation of true distances <50 m, matching observed discrepancies between GPS and RSSI (**Fig. 7C**). Comparisons of percent error showed that GPS consistently overestimated close-range proximity (<50 m) relative to true distances, with higher error than RSSI-based estimates from **Experiments 1–3** (**Fig. S22**). These results suggest a tradeoff in the accuracy of RSSI- and GPS-based inferences, with RSSI potentially out-performing GPS at pairwise distances <50 m.

## Discussion

Accurately quantifying spatial proximity is central to understanding social behavior, information flow, and pathogen transmission in animals (Krause et al. 2013, Smith and Pinter-Wollman 2021, Suraci et al. 2022). Here, we used a series of controlled experiments and field trials with multi-sensor WildFi biologgers (Wild et al. 2022b) to evaluate the utility of WiFi-based RSSI as a continuous proxy for proximity and localization. We found that RSSI can produce biologically useful estimates of distance, outperform GPS at close range, and support trilateration over narrow spatial extents. Across applications augmenting RSSI with tag-measured environmental data, other contextual data, and information about tag variation significantly benefited inferences.

RSSI was an effective proxy for pairwise distance, with conditional *R^2^* values of 0.771 - 0.936 across experiments. Out-of-sample cross-validations supported RSSI as an effective tool for predicting proximity in new contexts, with median error often <50% of the true distance and exceptional contact classification accuracy (>85-95%) for pairwise distances <2 m. However, our work also echoes previous findings, showing that RSSI is influenced by tag orientation, weather, obstructions, and tag variation (Prange et al. 2006, Böhm et al. 2009, Drewe et al. 2012, Boyland et al. 2013, Marfievici et al. 2013, Bettaney et al. 2015, Rutz et al. 2015). Building from previous work, we highlight how multi-sensor data can be used to refine RSSI-based inferences. Specifically, tag-measured temperature was universally favored by AIC and modestly increased *R^2^* by 1-3%. Our findings also reinforce the importance of context-specific calibration (Rutz et al. 2015), with 10-20% of variation in distance often resulting from tag random effects. While RSSI alone can serve as an effective proxy, pre-deployment validations and tag-measured data provide substantial performance gains with richer behavioral inference than RSSI alone.

Given the predictive accuracy of RSSI, we could trilaterate tag locations within a 1.3 ha gateway array with median Euclidean error of 2.6-3.3 m, accurate enough to reconstruct proximity networks. Including line-of-sight conditions, tag-measured temperature, and tag random effects further improved localization accuracy and network correlations. Compared to RFID arrays, which are limited to point detections, and GPS, which may be unavailable, too energy intensive, or too imprecise in some systems, RSSI trilateration offers a compelling middle ground – providing continuous spatial estimates over narrow spatial extents with lower noise and fewer outliers. While performance will vary with gateway layout and density (Kluge et al. 2020, Ripperger et al. 2020), these findings suggest that RSSI can make localization and indirect spatial interactions viable for behavioral inferences.

In free-ranging Egyptian fruit bats, RSSI was correlated with GPS-inferred distances but with greater variability than in controlled settings. While RSSI remained predictive across a broad range (up to 250 m), its relationship with GPS was nonlinear and noisier - partially reflecting positional uncertainty in the GPS data itself. Simulations using empirical GPS error confirmed that GPS systematically overestimated close-range distances (<50 m) as shown by He et al. (2023), consistent with divergence between RSSI- and GPS-based proximity networks at these scales. As a result, GPS and RSSI were weakly correlated (if at all) at distances <50 m, suggesting that GPS and RSSI differ most at the short ranges biologically relevant to epidemiological contact or direct social interaction. Interestingly, RSSI and GPS estimates were relatively well-correlated at intermediate-to-longer ranges (100-250 m), suggesting that some RSSI signals may also capture broader-scale spatial associations. There is increasing interest in the fine-scale interactions shaping animal societies and epidemiology through GPS tracking (Manlove et al. 2022, Wilber et al. 2022, He et al. 2023, Yang et al. 2023). Our findings highlight the importance of matching tools to the spatial and temporal scale of interest. For quantifying contacts or social structure within groups, RSSI may offer better resolution and lower bias than GPS. Used together, these technologies can provide complementary views of animal space use, proximity, and interaction across scales.

Our results demonstrate that RSSI is a practical and scalable tool for estimating fine-scale animal proximity and localization, particularly in conditions where GPS may fail. Rather than treating RSSI variability as noise, we show that environmental context and tag-specific effects can be leveraged to refine inference and move beyond binary contact definitions. This echoes calls for rigorous pre-deployment validation and context-specific calibration (Rutz et al. 2015), and we suggest that factors like ambient weather conditions should be considered when designing validation experiments. Importantly, our work also suggests that multi-sensor biologgers can help refine RSSI proximity sensing by capturing variation in ambient weather. Multi-sensor platforms are rapidly evolving and can accommodate many more data streams than weather alone (Williams et al. 2020). Including additional sensors (*e.g.*, accelerometers, magnetometers) on tags offers exciting opportunities to possibly further resolve behavioral effects on RSSI, accounting for the effects of motion, orientation, or other tag contexts. Synthesizing multiple data streams alongside RSSI can also offer avenues for edge computing (Yu et al. 2024) to create modeled “virtual sensors” of pairwise distance on tags that reduce data transmission costs and preserve biological inferences. More broadly, RSSI enables both accurate proximity sensing and trilateration, offering a powerful framework for high-resolution inference of animal interactions, advancing our ability to study social behavior in the wild.

## Acknowledgements

We thank Simon Amato for technical support and the University of Konstanz Biological Sciences M.Sc. Vertiefungskurs 2023 and 2024 cohorts for field assistance. W.R. was supported by a Gruber Science Foundation Fellowship, funds from the Yale-Max Planck Center, Yale Institute of Biospheric Sciences, and the Lindsay International Research Fellowship. HJW is supported by a VolkswagenStiftung Freigeist Fellowship Az-9B258. JEK is supported by the PRIME programme of the German Academic Exchange Service (DAAD) with funds from the German Federal Ministry of Education and Research (BMBF) and by the NOMIS Foundation. This research is funded in part by the Gordon and Betty Moore Foundation through Grant GBMF10539 and by the Academy for the Protection of Zoo Animals and Wildlife to MW and through the US National Institutes of Health through Grant # NIH 1R01GM131319 to VOE.

