## Supplementary Information for "Fine-scale animal proximity detection and localization via multi-sensor biologgers"

Contents:

Tables S1-4	Pages 2-5

Figures S1-22	Pages 6-27

**Table S1:** Out-of-sample model performance metrics of generalized linear mixed-effect models of known distance based on RSSI and orientation from Experiment 1 with Gamma log-link function (*n_tags_* = 2, *n_observations_* = 800). Performance metrics across 25 cross-validations are summarized using median and 95% intervals in parentheses.

| Model | Prediction Type | R^2^ | RMSE (m) |
| --- | --- | --- | --- |
| RSSI | Conditional | 0.503  (0.442, 0.53) | 3.87  (3.322, 4.876) |
|  | Mean | 0.503  (0.442, 0.53) | 3.87  (3.322, 4.876) |
| RSSI +  Orientation | Conditional | 0.833  (0.768, 0.859) | 1.356  (1.236, 1.717) |
|  | Mean | 0.834  (0.769, 0.859) | 1.356  (1.236, 1.719) |

**SI Table S2:** Out-of-sample model performance metrics of generalized linear mixed-effect models of known distance based on RSSI and tag-measured temperature from Experiment 2 with Gamma log-link function (*n_tags_* = 35, *n_observations_* = 269,443). Performance metrics across 25 cross-validations are summarized using median and 95% intervals in parentheses.

| Model | Prediction Type | R^2^ | RMSE (m) |
| --- | --- | --- | --- |
| RSSI | Conditional | 0.665  (0.661, 0.67) | 1.986  (1.969, 1.996) |
|  | Mean | 0.548  (0.544, 0.552) | 2.83  (2.798, 2.863) |
| RSSI +  Temperature | Conditional | 0.697  (0.693, 0.701) | 1.894  (1.877, 1.907) |
|  | Mean | 0.585  (0.582, 0.588) | 2.753  (2.721, 2.784) |

**Table S3:** Out-of-sample model performance metrics of generalized linear mixed-effect models of known distance based on RSSI-by-line-of-sight interactions and tag-measured temperature from Experiment 3 with Gamma log-link function (*n_tags_* = 17, *n_gateways_* = 16, *n_observations_* = 10,732). Performance metrics across 25 cross-validations are summarized using median and 95% intervals in parentheses.

| Model | Prediction Type | R^2^ | RMSE (m) |
| --- | --- | --- | --- |
| RSSI | Conditional | 0.794  (0.788, 0.802) | 3.78  (3.72, 3.859) |
|  | Mean | 0.516  (0.497, 0.54) | 5.922  (5.695, 6.053) |
| RSSI*Line of Sight + Temperature | Conditional | 0.838  (0.833, 0.843) | 3.281  (3.23, 3.331) |
|  | Mean | 0.578  (0.554, 0.605) | 5.765  (5.562, 5.937) |

**Table S4:** Out-of-sample model performance metrics of generalized linear mixed-effect models of GPS-inferred distance based on RSSI and tag-measured weather data from the Field Trial based on log-transformed RSSI with Gamma log-link function (*n_tags_* = 136, *n_gateways_* = 136, *n_observations_* = 2,948). Performance metrics across 25 cross-validations are summarized using median and 95% intervals in parentheses.

| Model | Prediction Type | R^2^ | RMSE (m) |
| --- | --- | --- | --- |
| RSSI | Conditional | 0.232  (0.159, 0.280) | 169.415  (143.868, 194.679) |
|  | Mean | 0.14  (0.102, 0.183) | 175.479  (151.172, 203.565) |
| RSSI + Weather | Conditional | 0.239  (0.159, 0.290) | 169.280  (144.651, 1194.064) |
|  | Mean | 0.154  (0.116, 0.194) | 175.479  (151.172, 203.565) |

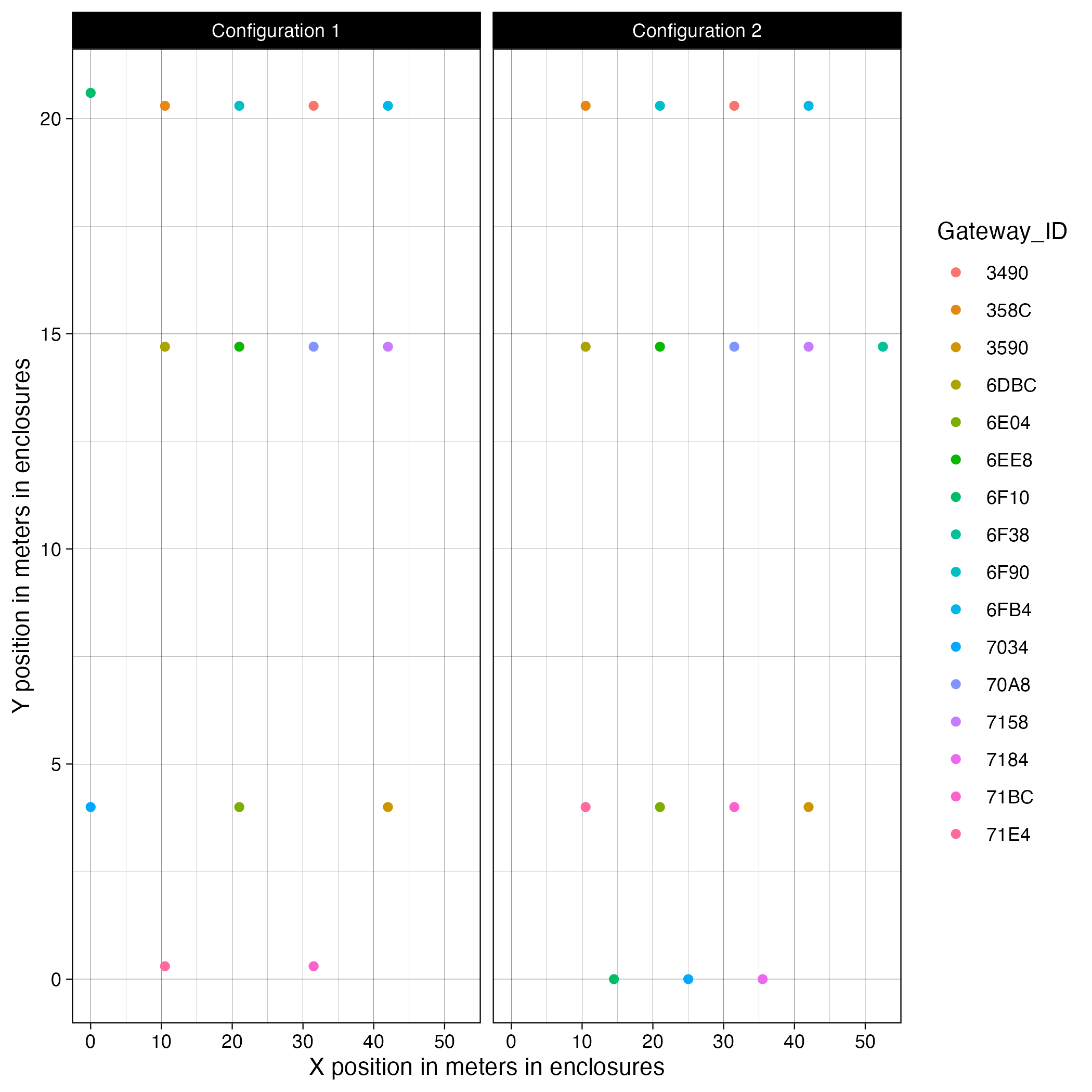

**Figure S1: Experiment 3 gateway layout**

Shown are the x and y coordinates of gateways within Experiment 3 (in meters). Configuration was treated as a random effect in analyses to address the shuffling of gateway locations between configurations.

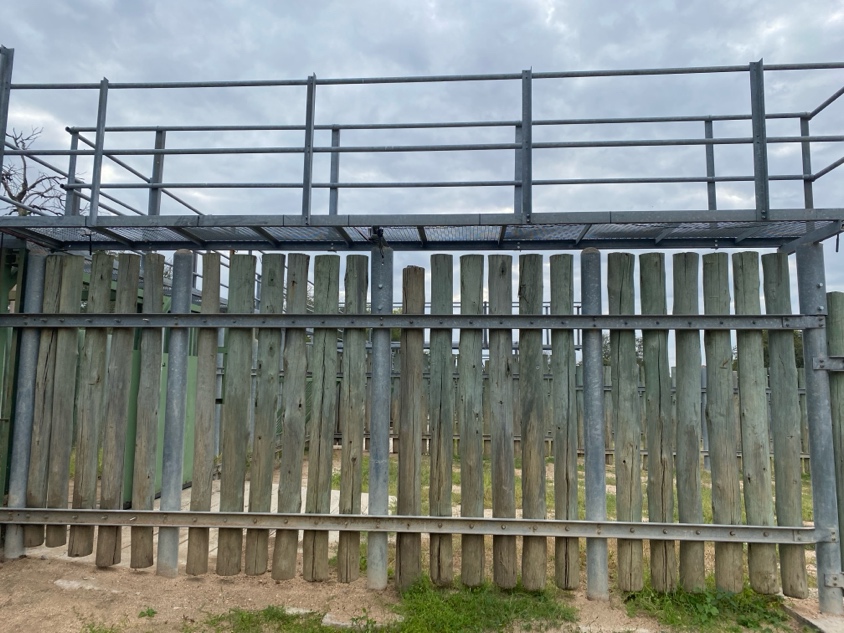

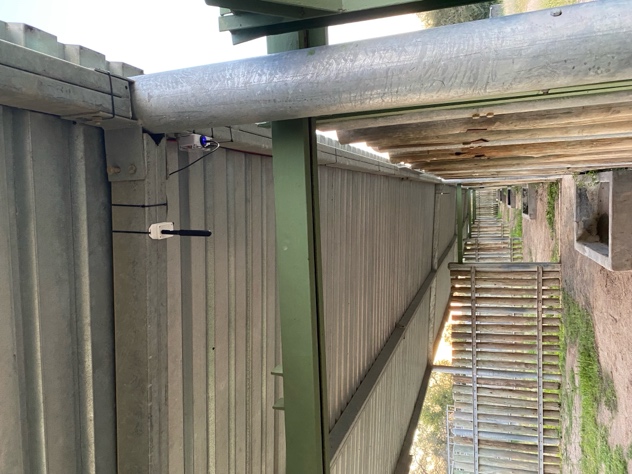

**Figure S2: Experiment 3 gateway position examples**

Photos of gateways (circled) positioned in enclosure setting: (left) depicts gateways positioned under a walkway; (right) depicts gateways underneath a shade covering.

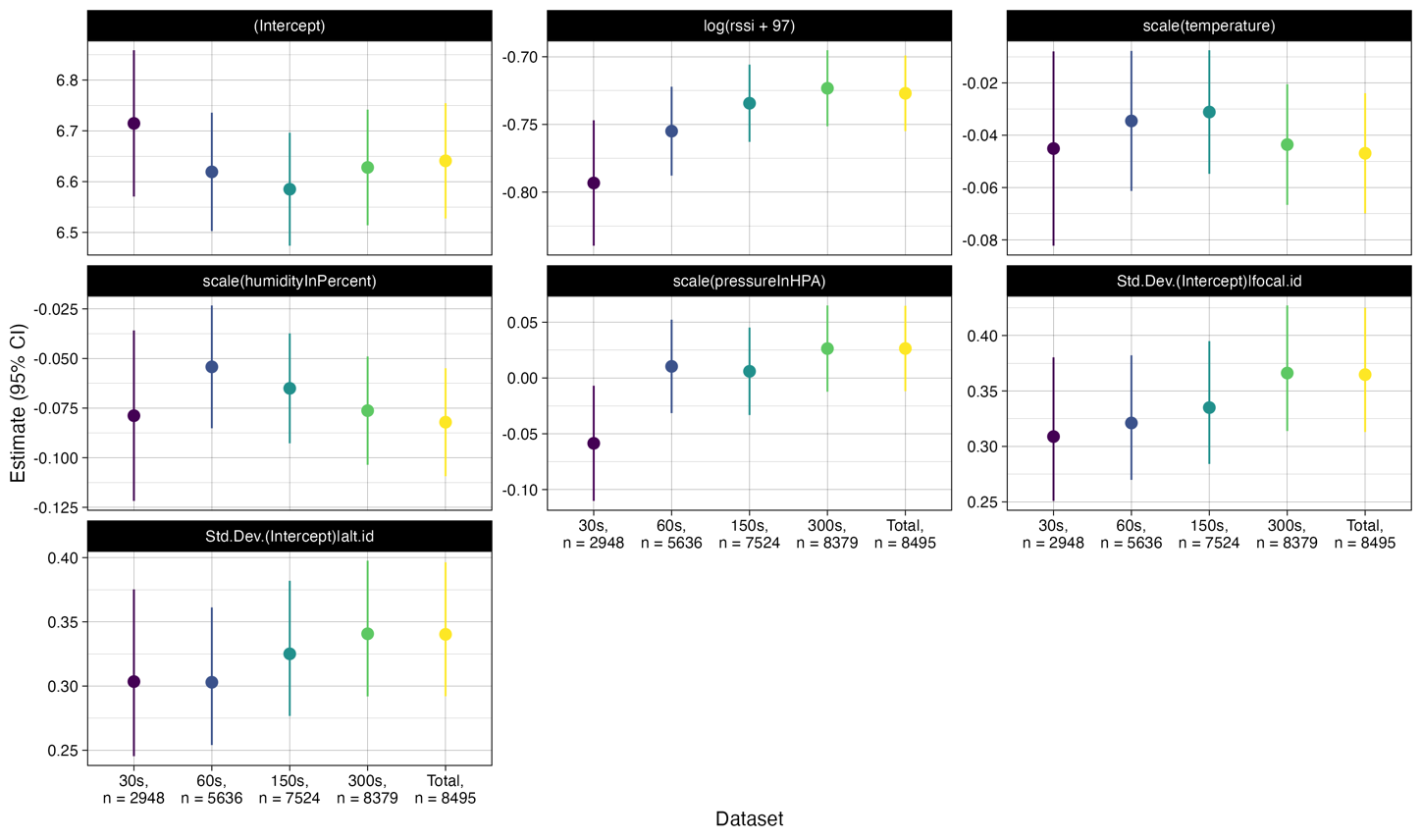

**Figure S3:** **Field Trial Approach 1 model coefficients based on time delay in GPS fixes**

Model estimates and 95% CI for coefficients and standard deviations of random effects under varying thresholds of total RSSI-focal GPS-alternate GPS time differences from 30s to 60s, 150s, 300s, and the entire, unfiltered dataset. Model estimates do differ significantly, but the size of these effects remains relatively small, indicating that GPS and RSSI asynchrony is impacting estimates but not overall patterns.

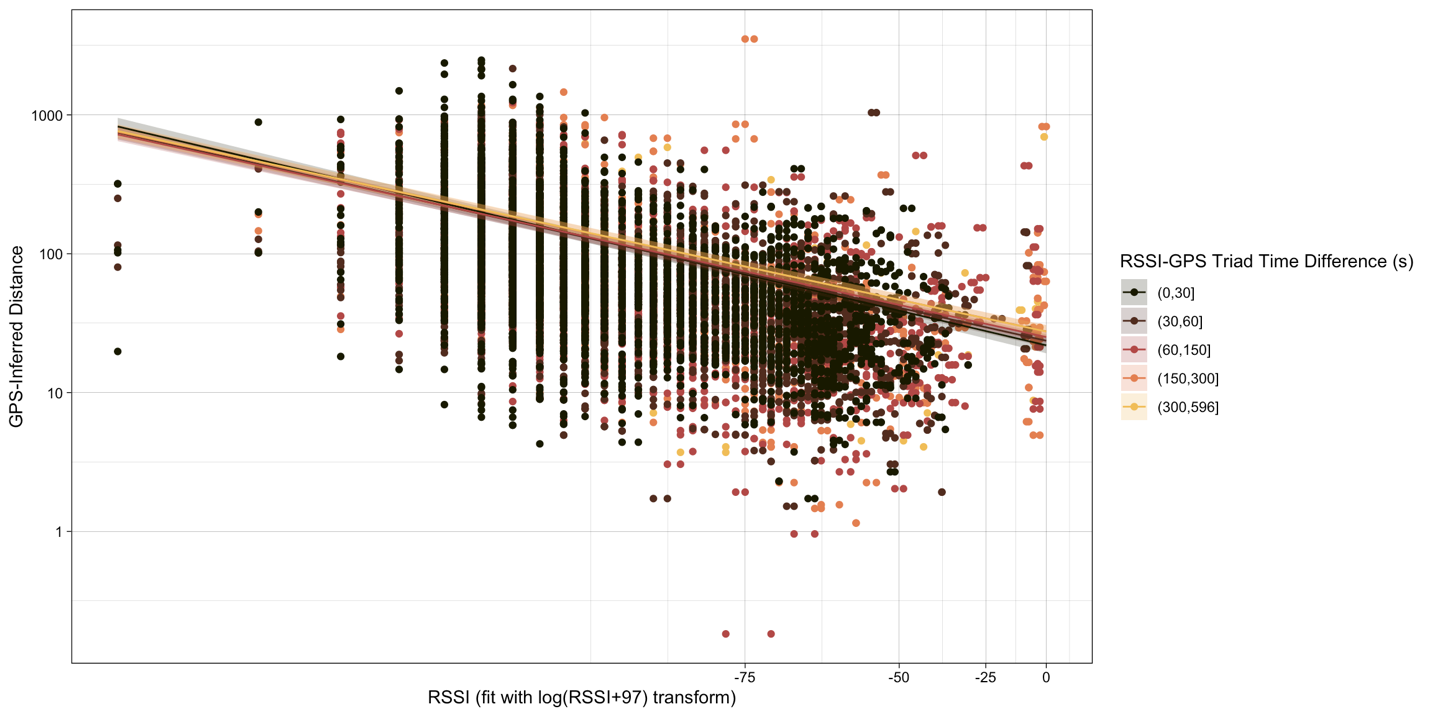

**Figure S4:** **Field Trial Approach 1 time delay in GPS fixes effect on RSSI relationship**

Depiction of the effect of total RSSI-focal GPS-alternate GPS time differences on inferred relationships between RSSI and GPS-inferred distances. Though there is variation in the estimates of models produced by different underlying datasets, the overall variation in these model estimates remains comparatively small relative to observed data.

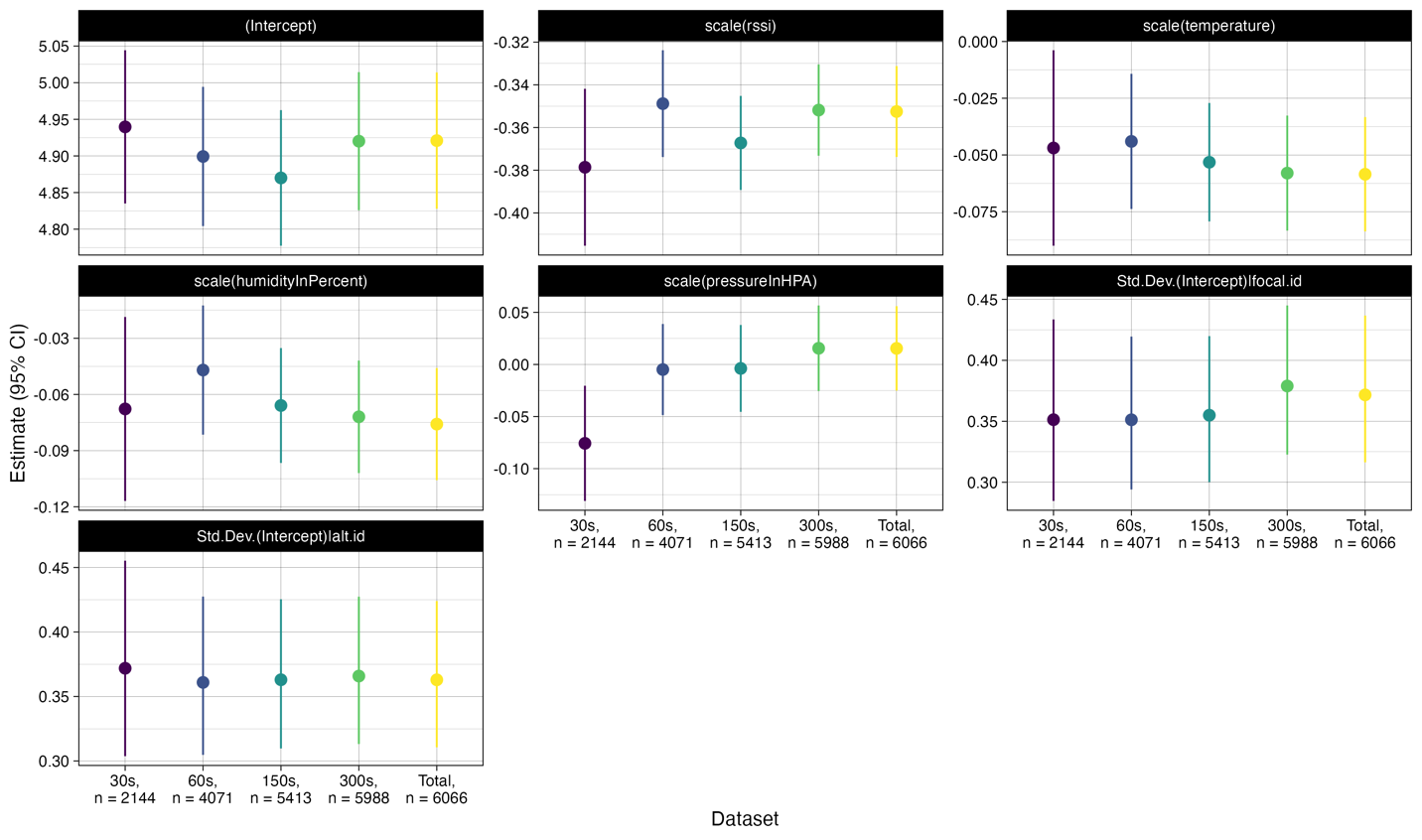

**Figure S5: Field Trial Approach 2 model coefficients based on time delay in GPS fixes**

Model estimates and 95% CI for coefficients and standard deviations of random effects under varying thresholds of total RSSI-focal GPS-alternate GPS time differences from 30s to 60s, 150s, 300s, and the entire, unfiltered dataset for RSSI values ≤-75. As for the modeling approach with log-transformed RSSI, the influence of RSSI and GPS asynchrony on inferences about the relationship between RSSI and distance remains small but statistically differentiable.

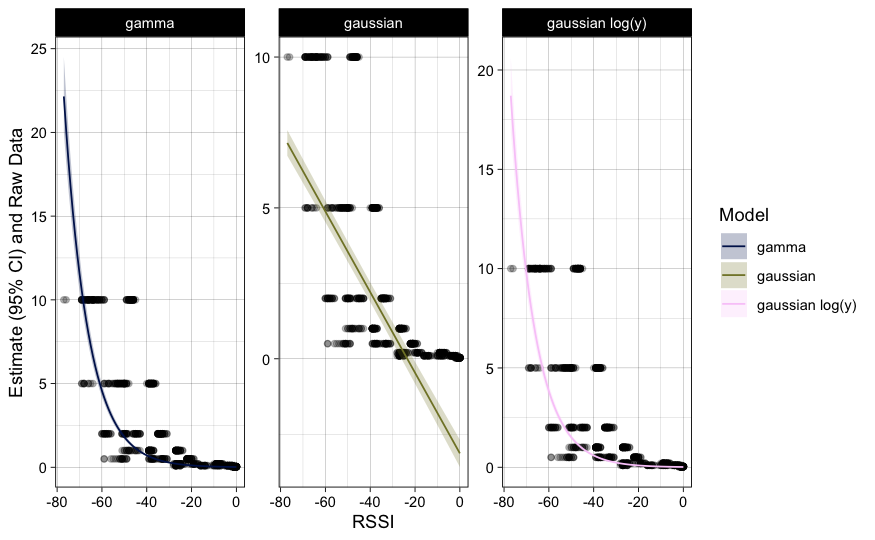

**Figure S6: Linear modeling Experiment 1 data with visual comparison**

Candidate link families for linear modeling under **Experiment 1** compared against raw data. Model predictions are based on the population mean (not conditioned on tag identity).

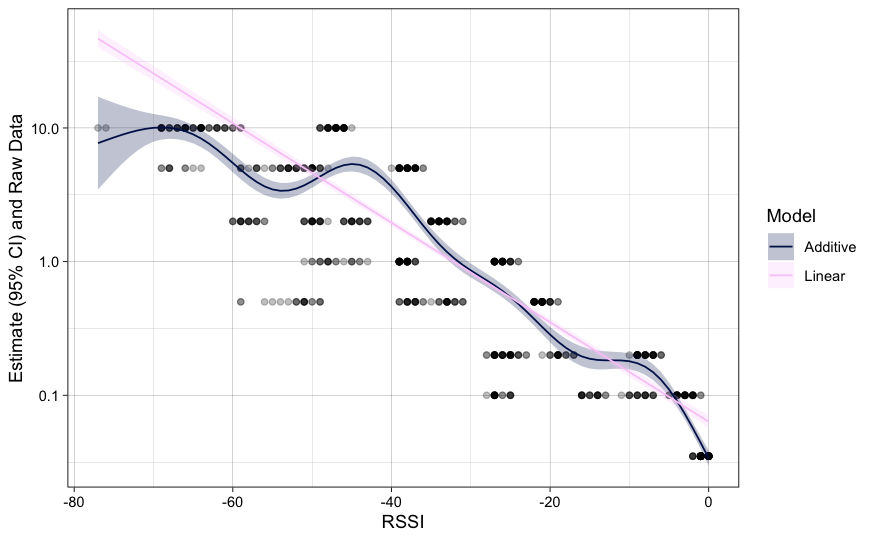

**Figure S7: Experiment 1 linear and additive model visual comparison**

Candidate linear and additive model fits under **Experiment 1** compared against raw data. Model predictions are based on the population mean (not conditioned on tag identity). Though there is more sinuosity in the additive fit, it visually appears that the linear model may adequately capture some of this pattern.

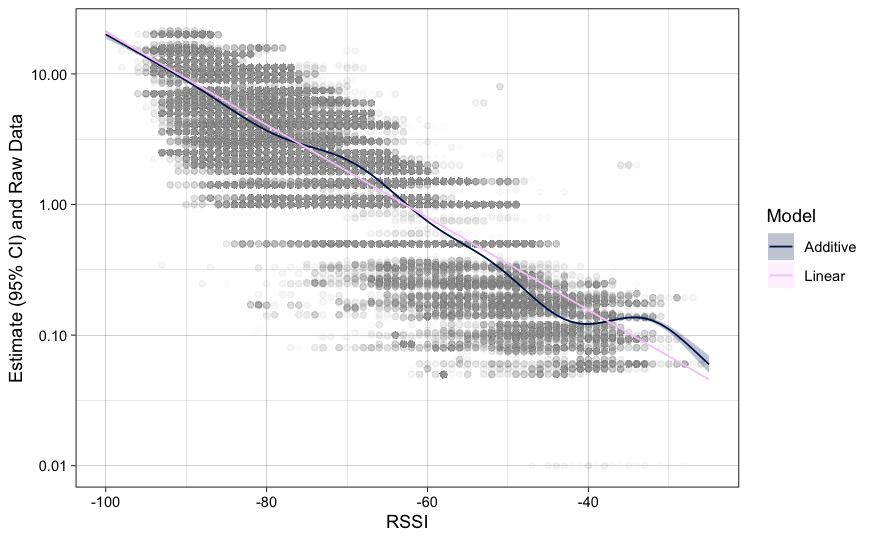

**Figure S8: Experiment 2 linear and additive model visual comparison**

Candidate linear and additive model fits under **Experiment 2** compared against raw data. Model predictions depicted here do not include random effects given the extreme complexity and dataset size for additive model fits. Though there is more sinuosity in the additive fit, it visually appears that the linear model may adequately capture some of this pattern.

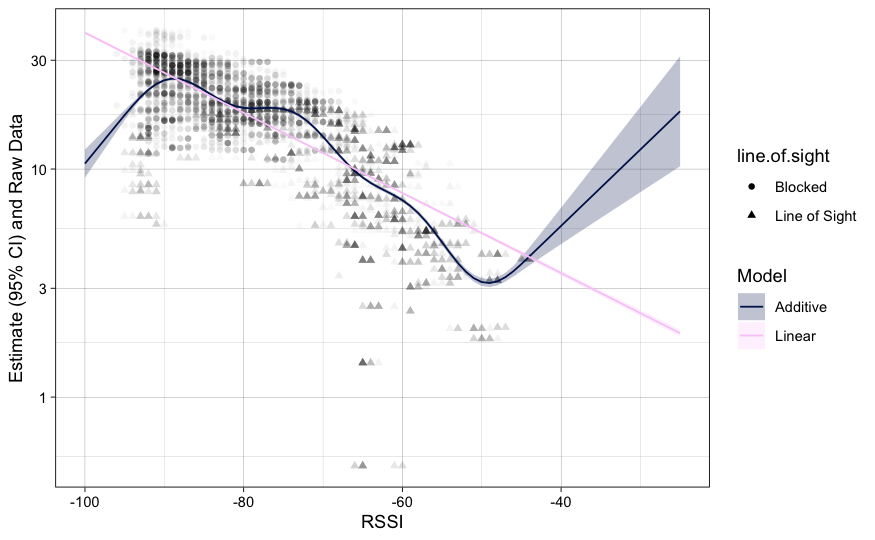

**Figure S9:** **Experiment 3 linear and additive model visual comparison**

Candidate linear and additive model fits under **Experiment 3** compared against raw data. Model predictions depicted here do not include random effects given the extreme complexity and dataset size for additive model fits. There is a distinct non-linearity in the additive model fit that is not captured in the linear model fit. This break largely corresponds to the segregation between tag-gateway communications that are blocked vs clear line-of-sight. This prompts the inclusion of an RSSI-by-line-of-sight interaction in later modeling.

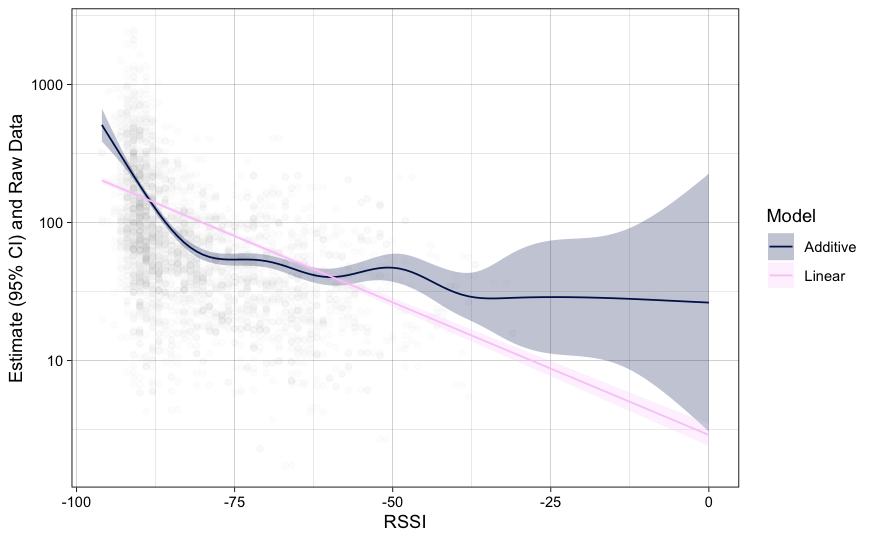

**Figure S10:** **Field Trial Approach 1 linear and additive model visual comparison**

Candidate linear and additive model fits under the **Field Trial** compared against raw data. Model predictions depicted here do not include random effects given the extreme complexity and dataset size for additive model fits. There is a distinct non-linearity in the additive model fit that is not captured in the linear model fit. This plateau in observed GPS data significantly affects linear model fit, prompting the inclusion of a log-RSSI term as well as a parallel modeling approach filtering based on RSSI values.

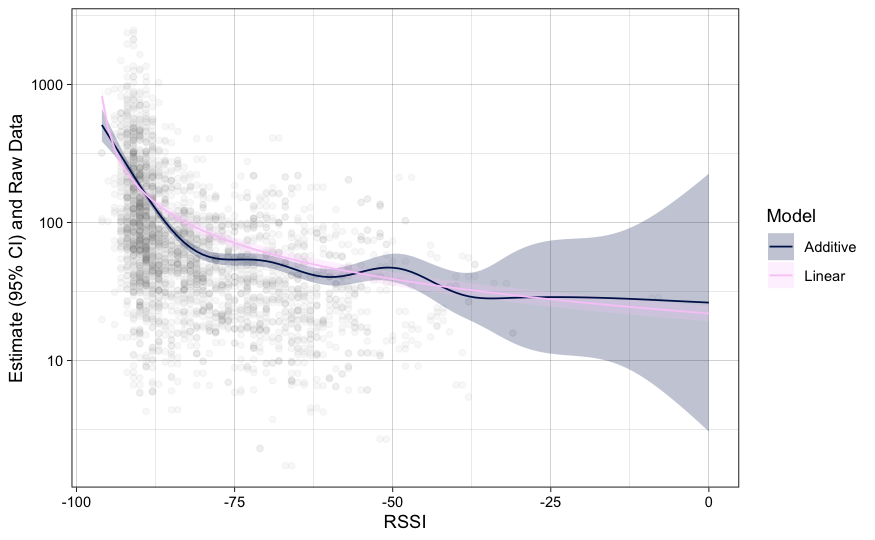

**Figure S11:** **Field Trial Approach 1 transformed linear and additive model visual comparison**

Updated candidate log-transformed RSSI linear and additive model fits under the **Field Trial** compared against raw data. Model predictions depicted here do not include random effects given the extreme complexity and dataset size for additive model fits. The inclusion of log-transformed RSSI values greatly reduced issues of fit between the linear and additive model. However, there remains a sharper decline in GPS distance vs RSSI values until about -80 to -75 RSSI. This prompts the filtering of the dataset to just these values to more accurately describe the linear change in RSSI relative to GPS as seen in **Experiments 1-3**.

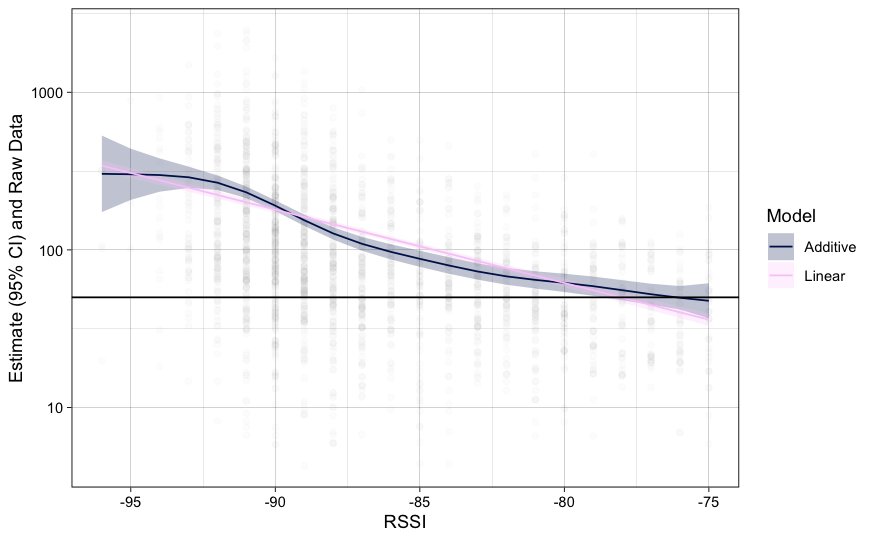

**Figure S12:** **Field Trial Approach 2 linear and additive model visual comparison**

Updated filtered RSSI linear and additive model fits under the **Field Trial** compared against raw data. Model predictions depicted here do not include random effects given the extreme complexity and dataset size for additive model fits. Subsetting the dataset greatly reduced the visual disparity between additive and linear model fits, and this approach maintains the same conceptual relationship with RSSI declining exponentially with distance as under **Experiments 1-3**.

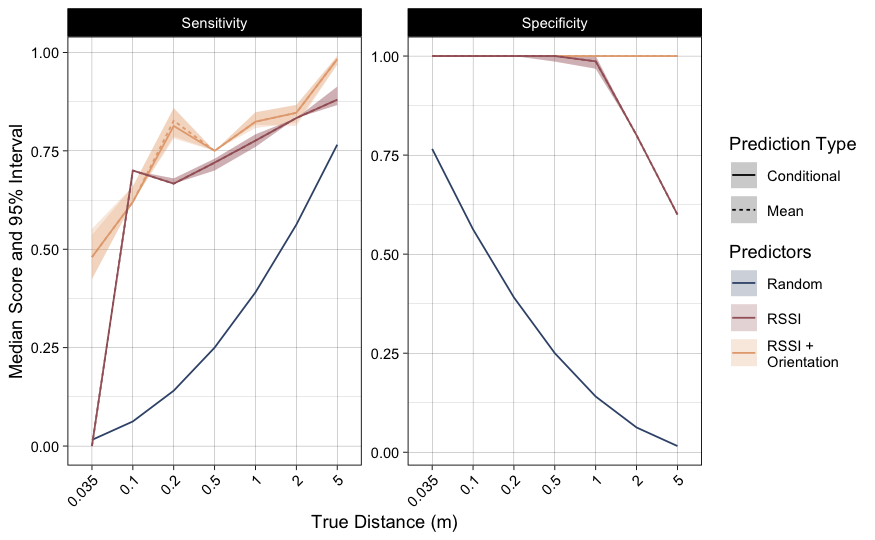

**Figure S13:** **Sensitivity and specificity of models in Experiment 1 cross-validations**

Median and 95% intervals for model sensitivity and specificity in classifying tag-tag communications at or below various true distance thresholds. Solid lines represent metrics based on predictions incorporating tag-level random effects, while dashed lines reflect population-only main effects. Metrics for a random classifier (blue) are also included for comparison, derived from the proportion of true contacts at each threshold.

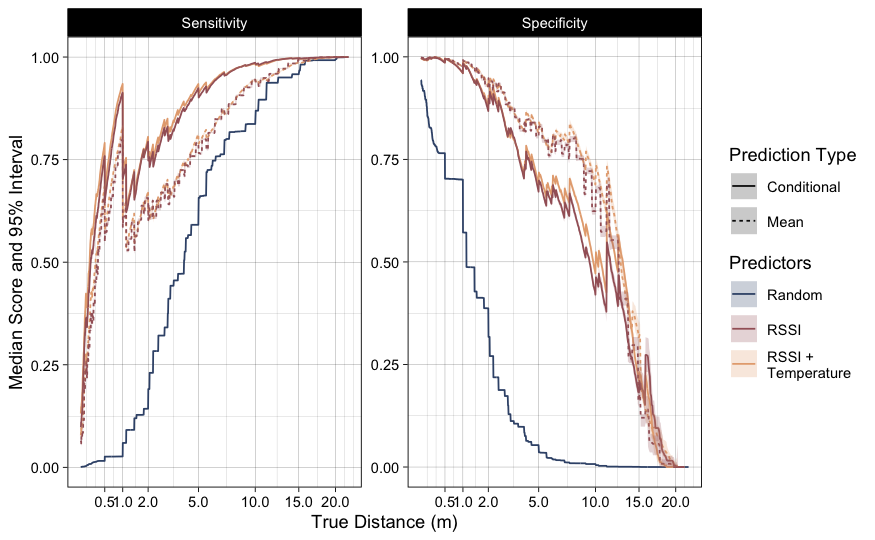

**Figure S14:** **Sensitivity and specificity of models in Experiment 2 cross-validations**

Median and 95% intervals for model sensitivity and specificity in classifying tag-tag communications at or below various true distance thresholds. Solid lines represent metrics based on predictions incorporating tag-level random effects, while dashed lines reflect population-only main effects. Metrics for a random classifier (blue) are also included for comparison, derived from the proportion of true contacts at each threshold.

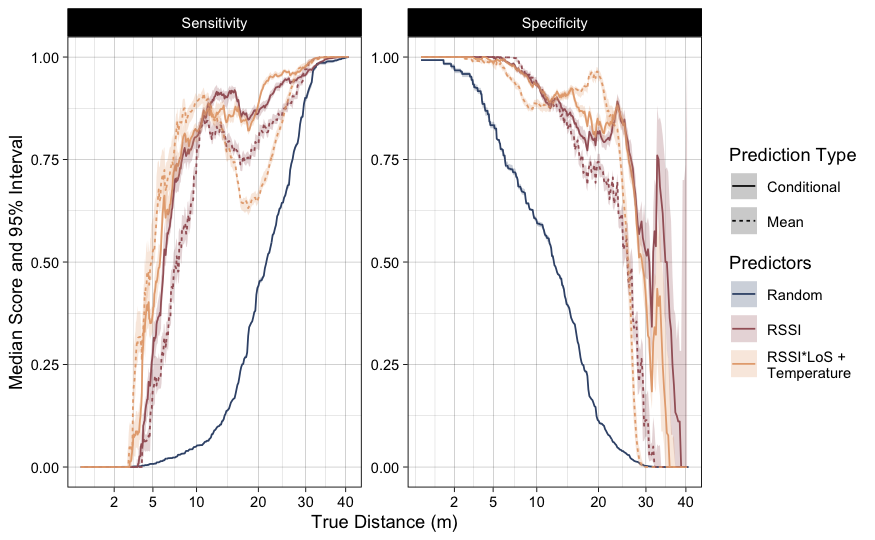

**Figure S15:** **Sensitivity and specificity of models in Experiment 3 cross-validations**

Median and 95% intervals for model sensitivity and specificity in classifying tag-tag communications at or below various true distance thresholds. Solid lines represent metrics based on predictions incorporating tag-level random effects, while dashed lines reflect population-only main effects. Metrics for a random classifier (blue) are also included for comparison, derived from the proportion of true contacts at each threshold.

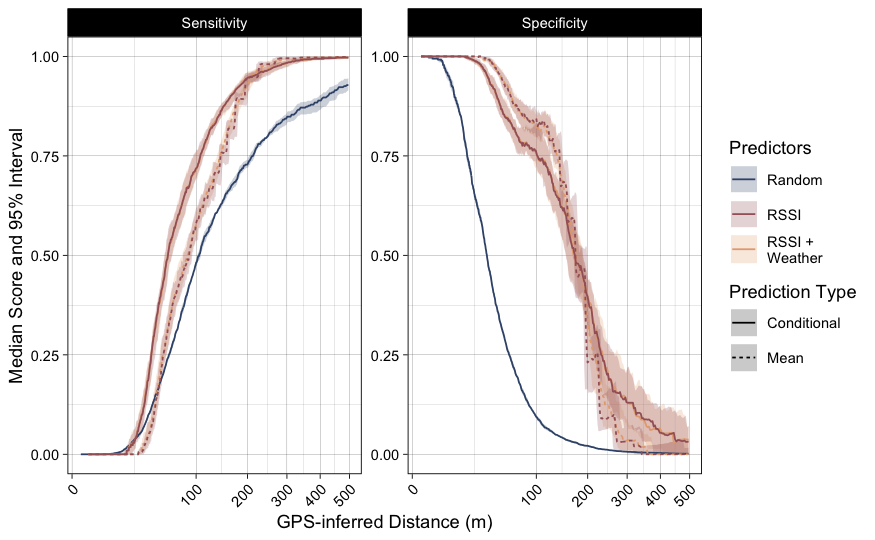

**Figure S16:** **Sensitivity and specificity of models in the Field Trial cross-validations**

Median and 95% intervals for model sensitivity and specificity in classifying tag-tag communications at or below various true distance thresholds. Solid lines represent metrics based on predictions incorporating tag-level random effects, while dashed lines reflect population-only main effects. Metrics for a random classifier (blue) are also included for comparison, derived from the proportion of true contacts at each threshold.

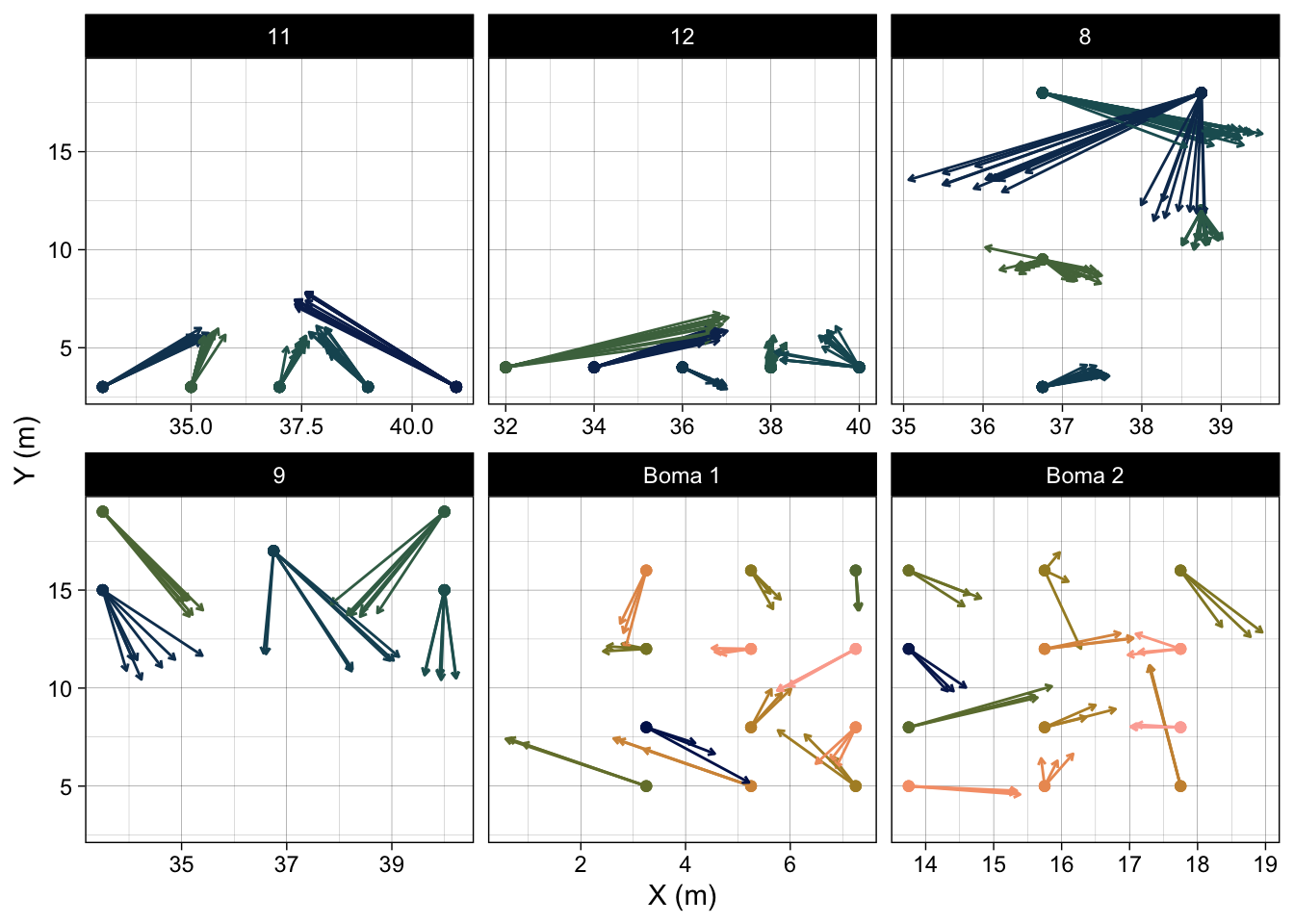

**Figure S17: Known and predicted tag locations from trilateration in Experiment 3**

Shown are the known coordinates of multi-sensor biologgers (points) and predictions from non-linear least squares regression models constructed from RSSI distance predictions, conditioned on line-of-sight, tag-measured temperature, and tag and gateway random effects (shown by arrows linked to true location). Facets denote different tag arrays within enclosure gateways.

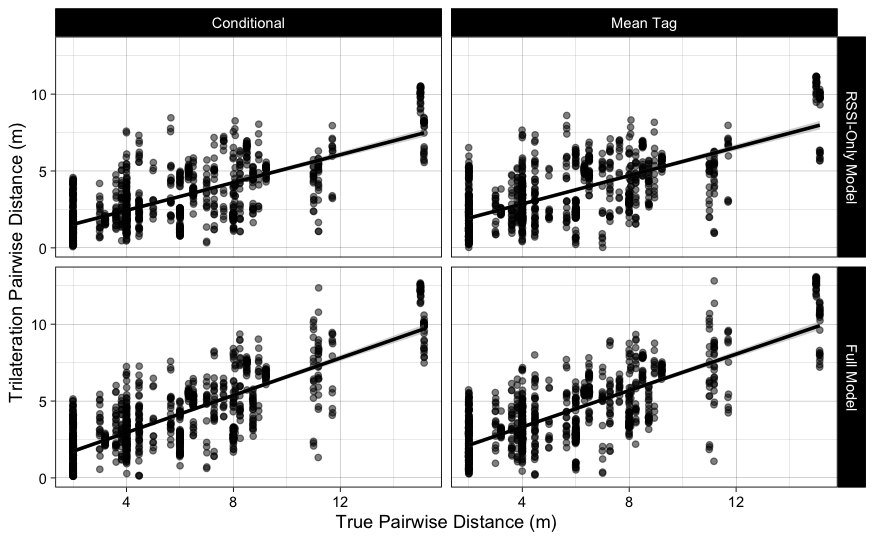

**Figure S18: Known and predicted pairwise distances from trilateration in Experiment 3**

Shown are the known pairwise distances of multi-sensor biologgers and pairwise distances between locations from trilateration predicted locations. Estimated distances are shown for a RSSI-only model and for a full model including a line-of-sight-by-RSSI interaction and tag-measured temperature. Predictions from RSSI-only and full models are shown either conditioned on tag random effects, or for a population-mean tag. Simple linear regressions between distances are shown for visual summary.

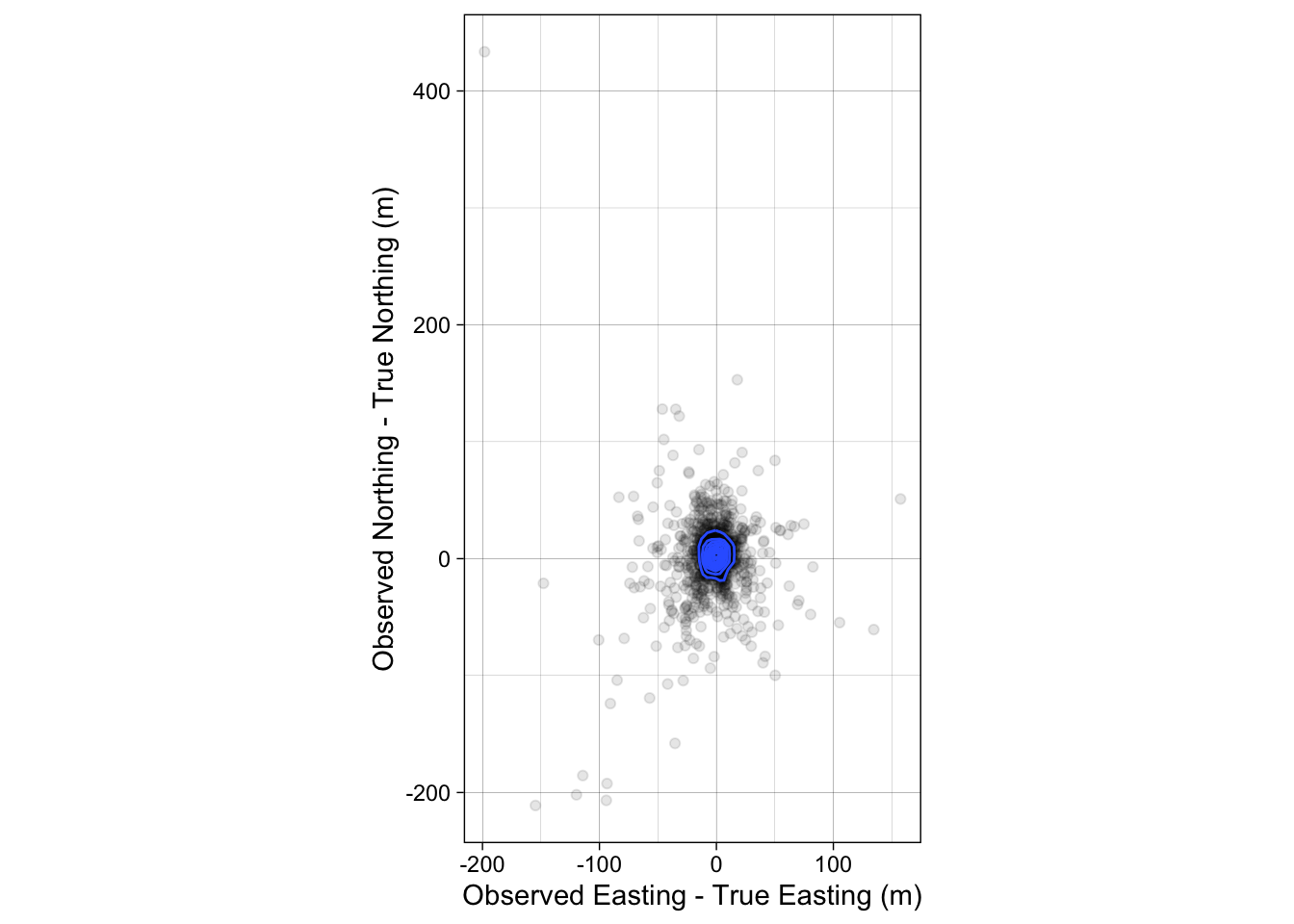
**Figure S19: Observed GPS error from stationary trial**

Stationary trial of the GPS chip used in the **Field Trial**. Easting and Northing are centered on the true location (*e.g.*, true x-y coordinates subtracted from observed). A density gradient is highlighted by blue contour lines.

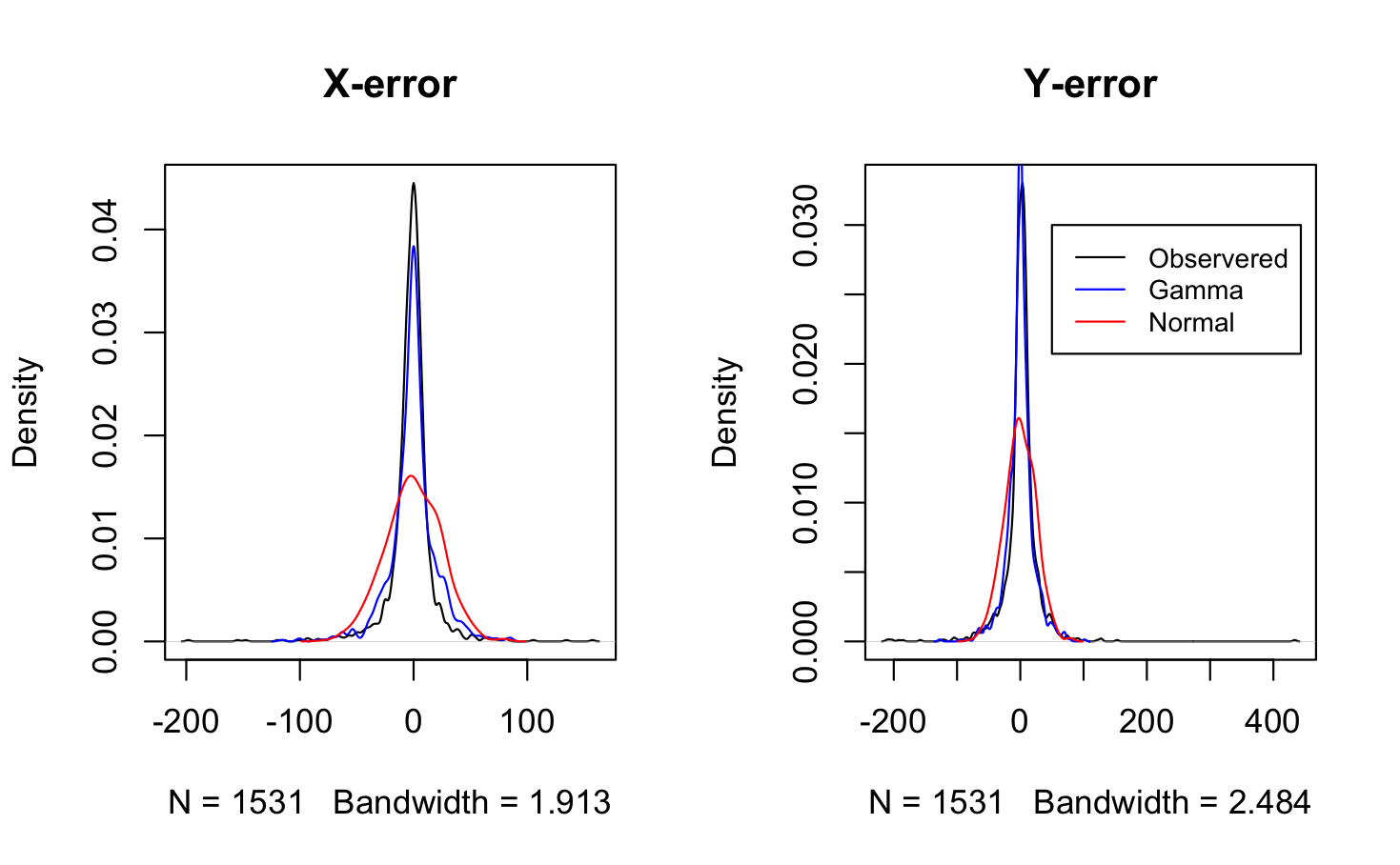

**Figure S20: Observed vs modeled GPS error in x and y dimensions**

Based on the stationary GPS trial, error in easting (x) and northing (y) in meters of the observed data (black) vs. modeled GPS error based on a normal distribution with sd equal to the observed sd of locations (red) or based on a gamma distribution parametrized from observed location error (blue). The observed error structure is more similar visually to the gamma distribution, with very high concentrations at 0 and heavier tails than present in the normal distribution.

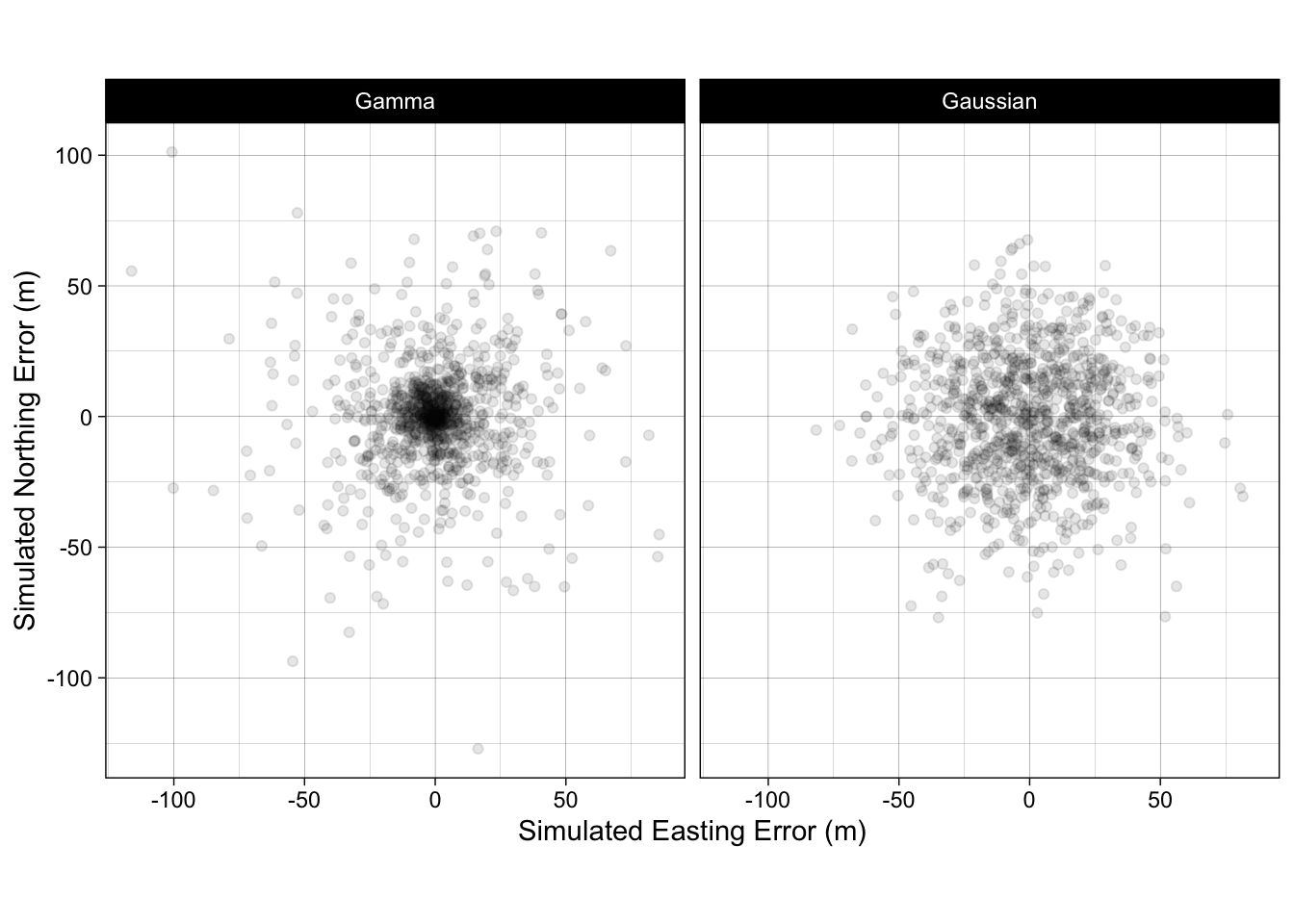

**Figure S21: Modeled GPS positional error in x and y dimensions**

Based on the stationary GPS trial, simulated GPS positions with error based on a normal distribution with sd equal to the observed sd of locations (right) or based on a gamma distribution parametrized from observed location error (left). The gamma, visually, appears to better match the patterns seen in **Figure S14**.

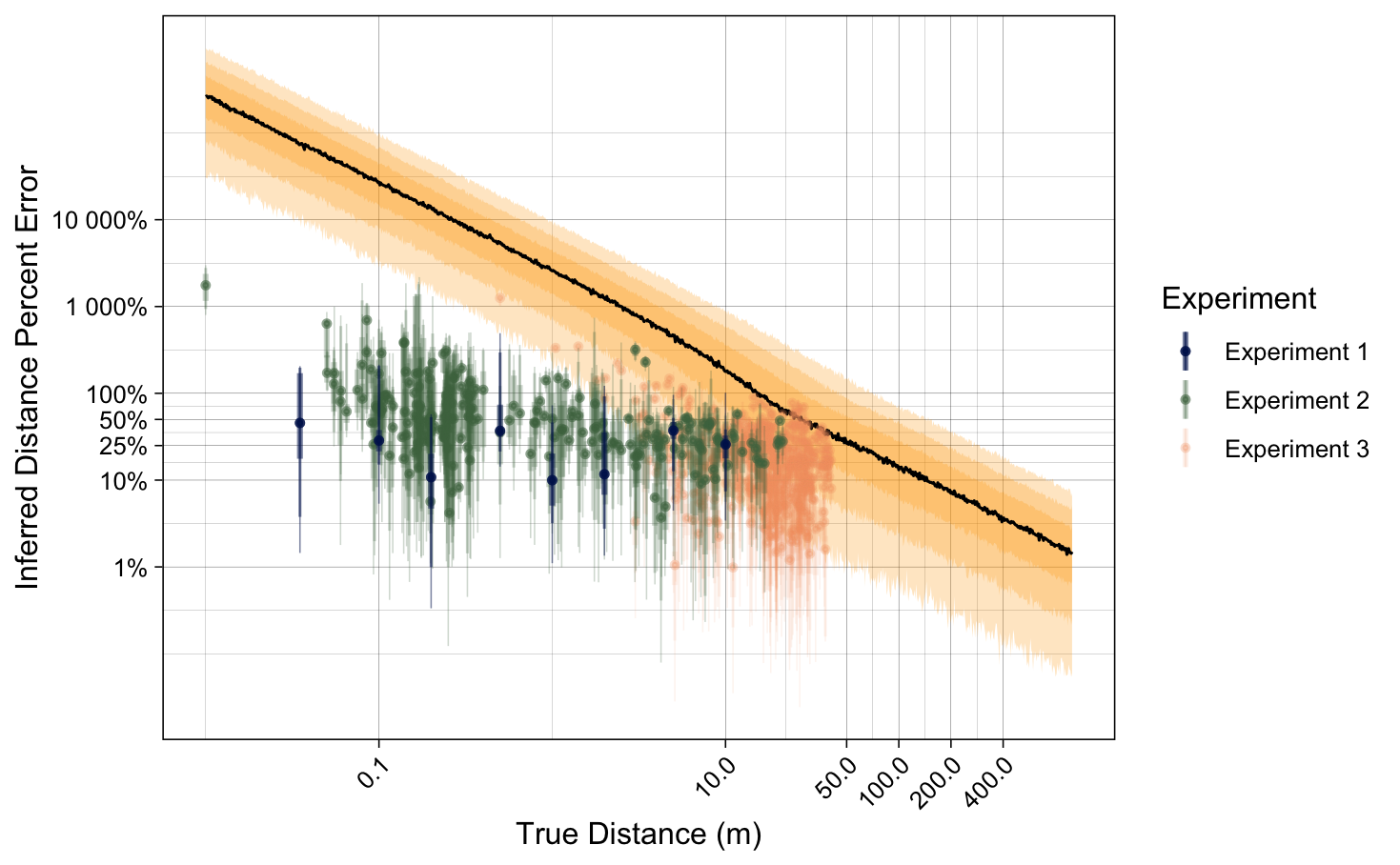

**Figure S22: Simulations of GPS vs. observed RSSI percent error**

Percent error in distance based on GPS estimates inclusive of positional error or RSSI model estimates based on data from **Experiments 1-3**. GPS estimates are based on pairwise distances between 1000 simulated point pairs, where location error was drawn from fitted gamma distributions. RSSI model estimates come from the top model for each experiment and are aggregated by unique true distances. For both estimate types, the median, 50%, 80%, and 95% intervals of percent error are shown.
